# Directed inter-domain motions enable the IsdH *Staphylococcus aureus* receptor to rapidly extract heme from human hemoglobin

**DOI:** 10.1101/2021.12.28.474365

**Authors:** Joseph Clayton, Kat Ellis-Guardiola, Brendan Mahoney, Jess Soule, Robert T. Clubb, Jeff Wereszczynski

## Abstract

Pathogenic *Staphylococcus aureus* actively acquires iron from human hemoglobin (Hb) using the IsdH surface receptor. Heme extraction is mediated by a tridomain unit within the receptor that contains its second (N2) and third (N3) NEAT domains joined by a helical linker domain. Extraction occurs within a dynamic complex, in which receptors engage each globin chain; the N2 domain tightly binds to Hb, while substantial inter-domain motions within the receptor enable its N3 domain to transiently distort the globin’s heme pocket. Using molecular simulations, Markov modeling, and quantitative measurements of heme transfer kinetics, we show that directed inter-domain motions within the receptor play a critical role in the extraction process. The directionality of N3 domain motion and the rate of heme extraction is controlled by amino acids within a short, flexible inter-domain tether that connects the N2 and linker domains. In the wild-type receptor directed motions originating from the tether enable the N3 domain to populate configurations capable of distorting Hb’s pocket, whereas mutant receptors containing altered tethers are less able to adopt these conformers and capture heme slowly via indirect processes in which Hb first releases heme into the solvent. Thus, our results show inter-domain motions within the IsdH receptor play a critical role in its ability to extract heme from Hb and highlight the importance of directed motions by the short, unstructured, amino acid sequence connecting the domains in controlling the directionality and magnitude of these functionally important motions.

## INTRODUCTION

*Staphylococcus aureus* is an opportunistic bacterial pathogen that causes a wide range of life-threatening illnesses such as pneumonia, meningitis, endocarditis, toxic shock syndrome, bacteremia, and septicemia [1–3]. During infections, *S. aureus* relies on iron as a nutrient, which it extracts from human hemoglobin (Hb) using the iron-regulated surface determinant (Isd) system [4–7]. In this process, Hb or its complex with human haptoglobin (Hp) is first captured on the bacterial surface by the homologous IsdB and IsdH receptors [8–12]. Both receptors actively extract Hb’s heme, which is then passed via cell wall associated hemoproteins (IsdA and IsdC) to the membrane-embedded ABC transporter complex (IsdEF) [13–15]. The heme is then pumped into the cytoplasm by the transporter, where it is degraded by oxygenases (IsdG and IsdI) to liberate its iron [16, 17]. The cell wall-associated proteins bind heme and Hb using NEAT (NEAr iron Transporter) domains, which are conserved in other Gram-positive bacteria [18, 19]. Understanding how heme is extracted from Hb could lead to novel treatments to combat lethal bacterial infections, as genetic elimination of genes encoding Isd proteins decreases *S. aureus* virulence, and other clinically important pathogens employ similar protein machinery to acquire heme during infections [13, 15, 20–23].

The IsdB and IsdH Hb receptors share significant primary sequence homology and extract heme using a conserved tri-domain unit in which two NEAT domains are connected by a helical linker domain (**Fig. 1A**) [9, 24, 25]. The NEAT domains in each tri-domain unit have distinct functions; the N-terminal domain binds to Hb, while the C-terminal domain captures ferric heme (hereafter called hemin). The tri-domain unit in IsdH is formed by the N2 and N3 NEAT domains joined by a linker domain (IsdH^N2N3^), while in IsdB the unit is formed from its N1 and N2 NEAT domains (IsdB^N1N2^). The extraction unit (hereafter called IsdH^N2N3^)was first discovered in IsdH and is formed by its N2 and N3 domains and an intervening linker domain [9]. The domains function synergistically and each must be part of the same polypeptide in order to rapidly acquire hemin from Hb [9]. A 4.2 Å resolution crystal structure of the IsdH^N2N3^:Hb complex revealed two binding interfaces: one between N2 and the A-helix of Hb that is located distal to Hb’s bound hemin molecule (called the N2-Hb interface), and a second binding surface between Hb’s hemin binding pocket and the receptor’s linker and N3 domains (called the LN3-Hb interface) (**Fig. 1A**) [26]. A subsequent 2.6 Å structure of the IsdH^N2N3^-Hb complex revealed deformation of Hb’s F-helix, which contains the hemin iron-coordinating axial His87 residue [12]. Subsequent structural studies of the IsdB-Hb complex revealed a similar mode of binding and F-helix distortion, suggesting that IsdB and IsdH extract Hb’s hemin using a related mechanism [27]. Interestingly, cellular studies suggest that IsdB is the primary Hb receptor in *S. aureus*, while IsdH may function to prevent Hp-mediated removal of Hb from the blood in addition to extracting its hemin [21, 28].

**Figure 1.**
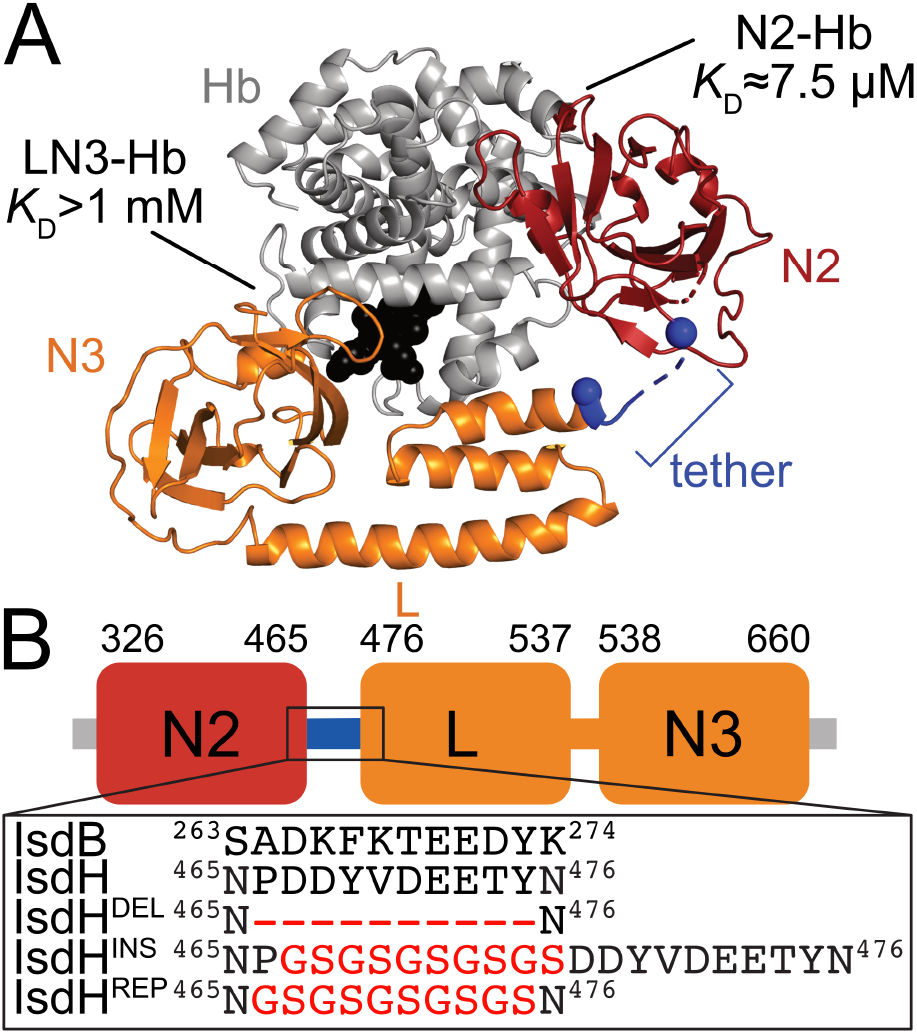
Structure of the IsdH-Hb complex. (A) Crystal structure of the IsdH:Hb complex (PDB code: 4XS0). The N2 (red) and LN3 (orange) subunits in the receptor form contacts with alpha globin chain in Hb (gray). The tether sequence and the N2-Hb and LN3-Hb interfaces are labeled. The hemin molecule in Hb is shown in space-filling format (black). The bounds of the tether (blue) are shown as spheres. (B) The panel shows the domain organization of IsdH with the amino acid sequence of the tether shown below. The tether sequence is formed by residues P466-Y475 (blue). Also shown are primary sequences of the tether mutants explored in this study, as well as the homologous tether sequence in IsdB.

Biophysical, computational, and structural studies of IsdH have shed light onto the mechanism underpinning hemin extraction. Hb binding is primarily mediated by the N2 domain, as the N2-Hb interface forms with a dissociation constant (K_D_) of 7.5 ± 0.1 µM that contributes ∼95% of the total binding standard free energy required for complex formation. In contrast, the LN3-Hb interface in which Hb’s F-helix is partially unwound forms with much weaker affinity (K_D_ > mM) [9, 11, 24]. Quantitative measurements that selectively measured the rate of hemin transfer from tetrameric Hb revealed that IsdH^N2N3^ accelerates hemin release from the alpha globin chain up to 13,400-fold as compared to the indirect process in which Hb first spontaneously releases hemin into the solvent, and identified two conserved receptor subsites within the Hb-LN3 interface that are important for transfer [9, 11]. Structural experiments and molecular dynamics (MD) simulations of the IsdH^N2N3^:Hb complex suggest that the receptor lowers the activation energy associated with hemin removal from the alpha subunit by inducing strain in the HisF8-Fe^3+^ axial bond and by increasing the concentration of nearby water molecules that compete with Hb’s HisF8’s Nε atom for the iron atom within hemin. After breakage of the HisF8-Fe^3+^ axial linkage the released hemin is then transferred ∼12 Å to the binding pocket in IsdH’s N3 domain where its metal is coordinated via the phenol oxygen of Y642 [11, 12, 26].

NMR studies indicate that the IsdH receptor undergoes significant inter-domain motions when bound to Hb [10, 29]. The motions occur at the junction between its N2 and linker domains, suggesting that LN3 domains form a rigid unit that transiently engages and distorts Hb’s hemin pocket while the N2 domain remains affixed to each globin [29]. Here, we show using computational and experimental approaches that these inter-domain motions are functionally important and defined by a semi-disordered tether sequence that connects the N2 and linker domains (residues P466-Y475). MD simulations analyzed through Markov state models of wild-type and mutant receptors reveal that the tether plays a key role in controlling both the positioning and direction of motion of the LN3 unit relative to Hb. These directed domain motions raise the effective molar concentration of the LN3 unit near Hb to ∼15 mM, enabling it to populate a low affinity Hb-LN3 interface in which Hb’s heme pocket is distorted to promote hemin release. Residues in the tether are mobile in the simulations and absent in crystal structures of Hb-IsdH complexes, yet nevertheless direct inter-domain motions by transiently adopting a closed conformation that projects the LN3 unit toward Hb. Altering the sequence of the tether disrupts its ability to direct functionally important domain motions dramatically slowing the rate at which hemin is captured.

## METHODS

### Molecular dynamics simulations

Initial atom coordinates for the WT IsdH were taken from a 2.55 Å resolution IsdH:Hb dimer structure (PDB: 4XS0) [12]; the hemoglobin dimer was deleted, and missing residues were built. The protein was then neutralized with Na+ and solvated with a 150 mM NaCl buffer using Joung/Cheatam monovalent ion parameters and TIP3P water [30, 31]. The protein was parameterized using the AMBER ff14SB [32] while the heme and Fe-N bond parameters were taken from previous work [29]. The system was minimized and heated to 300 K over 20 ps while restraining the protein, then the restraints were relaxed over 100 ps. Three separate 500 ns simulations were carried out under constant temperature and pressure with a Langavin thermostat with a collision frequency of 1 ps^-1^ and a Monte Carlo barostat set to 1 atm [33, 34]; electrostatic energies were calculated using particle-mesh Ewald while van der Waals energies were calculated with a cutoff of 10 Å. All simulations were done using the AMBER18 software suite with a timestep of 2 fs, unless otherwise specified [35]. Standard simulation analyses were performed with CPPTRAJ [36].

To further sample the interdomain motions, the N2-N3 distance and N2-H3-N3 dihedral was calculated for every 100 ps, then frames from the three simulations were clustered into 50 groups using these two projections. One structure from each group was then selected at random to seed a 100 ns simulation, where each seeding configuration was prepared as described above. Additional sampling was conducted using the goal-oriented FAST protocol [37]; the trajectory set was used to create a temporary Markov model based on the distance and dihedral projections. Clusters in the model were selected based on a reward function which used their proximity to the goal state and on the sampling of similar structures. Here, the goal state was any structure that was different than the initial crystal coordinates–that is, states with similar coordinates as the WT IsdH:Hb dimer structures were penalized. Selected structures were prepared and run for 100 ns, then added to the trajectory set. 20 rounds of this sampling were produced, leading to a total of 25 μs of sampling. The resulting trajectories from the FAST protocol were then represented by a final Markov model consisting of 200 states. All models were built using the KMeans clustering algorithm and a lag time of 50 ns was selected based off the implied timescales of the model. Distance and dihedral measurements were done using CPPTRAJ [36], while Markov model building and coarse-graining were done with MSMBuilder [38]. Visualization was done using PyMol version 2.3.0 [39].

To sample the N3 binding process, we first formed a coarse Markov model by lumping the 200 microstates into 75 macrostates using the BACE algorithm [40]. This algorithm was selected in order to clump clusters that behaved similar in the model and to ensure these clumps were sufficiently different. A representative apo-IsdH structure was taken from each macrostate, then aligned the N2 domain to the IsdH:Hb dimer structure (PDB:4XS0); the IsdHN2N3 in the crystal structure was then deleted, leaving the Hb structure and the aligned IsdHN2N3. Of these 75 initial structures, 51 did not lead to structural overlap between the N3 and Hb dimer. These structures were prepared as described above and simulated for 100 ns. This set was then used to initialized another FAST protocol of five rounds, where structures were rewarded for being similar to the WT IsdH:Hb dimer. The N3 RMSD relative to its positioning in the Hb bound crystal structure (after aligning the α-subunits) was included as a third dimension during the clustering. Outside the clustering dimensions selection, the protocol followed was the same as mentioned above.

Neither the initial simulations nor the FAST protocol were able to sample states similar to the N3:Hb interface seen in the crystal structure, thus the five aligned structures with the lowest N3 RMSD were used to initialize steered MD simulations where the N3 RMSD was steered towards zero over 50 ns with a force constant of 1.0 kcal/mol/Å. Initial structures where the N3 RMSD was 5, 10, and 15 Å were pulled from these trajectories and used to seed three conventional, unbiased 100 ns simulations–leading to 4.5 µs of sampling states similar to IsdH:Hb dimer structure. The steered simulations were done using NAMD2.12 [41] through the Colvars module [42], while the seeded simulations were conducted as described above.

IsdH-Hb trajectories from the initial aligned structures, the FAST rounds, and seeded from the steered simulations were then analyzed by building a Markov model for each structural mutation. The construction of the model was done in the same manner as the apo-IsdH models, except with the N3 RMSD included as a third dimension in the clustering and a lag time of 85 ns was used. Binding kinetics were analyzed by coarse-graining the model to 25 macrostates using BACE. Each structural mutation was built using Robetta [43] and sampled in the same way described above. Two separate mutations were introduced: a ten residue insertion consisting of an alternating Gly-Ser sequence in between Pro466 and Asp467, and a ten residue deletion (residues Pro466-Y475). Apo- and holo-IsdH mutant models were produced following the protocol described above.

### Preparation of mutant plasmids

All mutants are derived from the parent gene encoding “wild-type” ^α^IsdH^N2N3^ as an insert at BamHI(5’)-XhoI(3’) restriction sites the pET28a-derived plasmid pSUMO, which encodes an N-terminal small-ubiquitin-like modifier (SUMO) fusion and an N-terminal hexahistidine tag. Mutations to this insert were introduced through QuikChange mutagenesis and single-overlap extension (SOE) PCR using primers supplied in the supplementary information. When SOE was used, the agarose-gel purified assembled insert and plasmid backbone were digested with BamHI and XhoI (NEB) and the insert ligated with T4 ligase. Plasmid DNA was transformed into chemically competent BL21(DE3) *E. coli* cells and plated on Luria broth (LB)-agar supplemented with 50 µg/mL kanamycin sulfate. Colonies were harvested, propagated for sequencing, and stored as stocks at −80°C in 1:1 LB:50% glycerol supplemented with 25 µg/mL kanamycin sulfate.

### Preparation of IsdH and Hb

Mutants of apo-IsdH and apo-Mb(H64Y/V68F) were prepared as previously reported [44–46]. Briefly, 10 mL LB supplemented with 50 µg/mL kanamycin sulfate (LB/kan) was inoculated with *E. coli* BL21(DE3) cells bearing mutant pSUMO-IsdH or pSUMO-apoMb(H64Y/V68F) plasmid and grown overnight at 37°C with 200 rpm shaking. 2 L of LB/kan were inoculated with overnight culture and grown to OD600 = 0.6 with 200 rpm shaking at 37°C. Cultures were induced with 2 mL 1 M IPTG and incubated with 200 rpm shaking at 37°C for 20-24 hours. Cells were harvested by centrifugation at 7000 rpm for 10 minutes in 500 mL plastic collection vessels in a Beckman JA-10 centrifuge rotor. Cell pellets from all mutants were slightly brown in color indicative E. coli-sourced heme bound to IsdH. Cells were resuspended in 50-60 mL lysis buffer (50 mM NaHPO4, 300 mM NaCl, 10 mM imidazole, pH 7.4) and frozen overnight at −20°C. Thawed cell pellets were supplemented with Protease Inhibitor Cocktail, lysed by sonication, and clarified at 15,000 rpm in a Beckman JA-20 centrifuge rotor. Clarified lysates were applied to 5 mL HisPur Ni-NTA resin by co-incubation at 4°C for 20 minutes. Flowthrough was eluted and bound protein washed with 50 mL lysis buffer. Protein was denatured on-bead with 50 mL denaturing buffer (50 mM NaHPO4, 300 mM NaCl, 10 mM imidazole, 6 M guanidine hydrochloride, pH 7.4) to release bound heme [46]. Heme was washed away with 4 × 50 mL washes of a 1:4 v/v mixture of ethanol:denaturing buffer. The denatured apo-protein was refolded by an additional wash with 50 mL lysis buffer. Resin was resuspended in ∼30 mL lysis buffer and 1 mL Ulp1-SUMO protease added as a 1 mg/mL solution in 50% glycerol/lysis buffer. Resin mixture was incubated overnight with gentle rocking at 4°C to liberate IsdH. The protein was then eluted and resin washed with 2 × 30 mL lysis buffer, concentrated and buffer exchanged into heme transfer-sucrose buffer (20 mM NaHPO4, 150 mM NaCl, 0.45 M sucrose, pH 7.5) using 15 mL Amicon 10 kDa cutoff centrifugal filters. metHb0.1 was prepared as reported previously, with small modifications. Briefly, 50 mL LB supplemented with 10 µg/mL tetracycline (LB/tet) was inoculated with *E. coli* BL21(DE3) cells bearing plasmid pSGE1.1-E4, which encodes the tetramer-stabilized Hb mutant Hb0.1. Cultures were grown overnight at 37°C with 200 rpm shaking. 2 L of M9 minimal medium supplemented with 10 µg/mL tetracycline was prepared and inoculated with overnight culture and incubated at 37°C with 200 rpm shaking to OD600 = 0.4-0.6. Hb0.1 expression was induced by addition of 2 mL 8 mM hemin in 0.1 M NaOH and 2 mL 1 M IPTG. Expression was conducted at 25°C for 20-24 h with 200 rpm shaking. Cells were harvested by centrifugation at 7000 rpm for 10 minutes in 500 mL plastic collection vessels using a Beckman JA-10 centrifuge rotor. Cells were washed twice with Hb0.1 lysis buffer (20 mM Tris, 17 mM NaCl, pH 8.5) to remove excess heme, resuspended in 50-60 mL Hb0.1 lysis buffer and frozen overnight at −20C. Thawed cell mixtures were bubbled with carbon monoxide to prevent auto-oxidation. Cells were treated with 1 mL 1 M benzamidine HCl and lysed by sonication. The crude lysate was clarified at 15,000 rpm in a Beckman JA-20 centrifuge rotor. Clarified lysate was applied to 10 mL HisPur Co-NTA resin and incubated with gentle shaking at 4°C for 20 minutes to maximize binding. For purification, all buffers were sparged thoroughly with CO to ensure the Hb0.1 Fe(II)-CO state is maintained. The binding supernatant was eluted and resin washed with 50 mL Hb0.1 lysis buffer, 50 mL wash buffer 1 (20 mM Tris, 500 mM NaCl, pH 8.5), 50 mL wash buffer 2 (20 mM Tris pH 8.5), and finally eluted with 3 × 25 mL elution buffer (20 mM Tris, 100 mM imidazole, pH 8.5). The eluted fraction was applied to 5 mL Q-Sepharose resin equilibrated with 20 mM Tris pH 8.5. Bound Hb0.1 was washed with 50 mL 20 mM Tris pH 8.5 and eluted with 3 × 20 mL 30 mM NaHPO4 pH 6.9. Eluted fraction of Hb0.1-CO was concentrated to 500 µM concentration and oxidized with 5 molar equivalents of potassium ferricyanide (K_3_Fe(CN)_6_) for 1 hour at 4ºC. Excess oxidant was removed by gel filtration over Sephadex G-25, and eluted metHb0.1 was buffer exchanged into heme transfer-sucrose buffer as described for IsdH using 15 mL Amicon 3 kDa cutoff centrifugal filters.

### Hemin transfer measurements

Rates of hemin transfer were measured with an Applied Photophysics SX-20 stopped-flow spectrophotometer under the following conditions: 5 µM metHb0.1 (heme basis) and 150 µM heme acceptor (apo-IsdH or apo-Mb(H64Y/V68F)) in 20 mM NaHPO4, 150 mM NaCl, 0.45 M sucrose, pH 7.5. Temperatures were varied from 15-37°C. Each mutant tested was measured in technical triplicate with 1000-second acquisition time at 405 nm (IsdH trials) and 600 nm (apo-Mb(H64Y/V68F) trials). Rate constants k_fast_ and k_slow_ were obtained through fitting the Abs_405_ curve. The absorbance of the reaction mixture at 405 nm at any given time is the sum of absorbance contributions from holo-Hb0.1, holo-IsdH, and background absorbance (given by constant 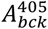).

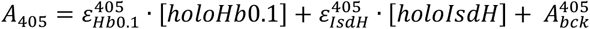

The transfer of hemin from Hb0.1 to IsdH results in an overall reduction in absorbance because 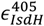 is lower than 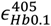. This process is fitted to a two-exponential decay function a fast phase and a slow phase, governed by rate constants k_fast_ and k_slow_. We constrain each process to 50% of the span, where span = *A*_405_(*t* = 0) − plateau.

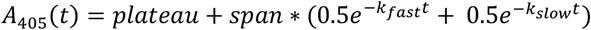

With limited experiment time, the plateau value for a given reaction is approximated as the minimum absorbance value obtained after reaction for 1000 seconds, the typical length of data acquisition. For fast mutants, like WT IsdH, this timeframe is sufficient to observe complete reaction with metHb0.1. Slower mutants do not react completely within 1000 seconds, so the span value is only a fraction of that observed with WT. In our analysis, the WT span is set as the maximum, describing complete transfer of 5 µM hemin from Hb0.1 to IsdH. For each slow mutant, we convert fractional amplitudes obtained after 1000 seconds to the concentration of hemin transferred as calibrated by the WT span. This allows us to report the rates in terms of the concentration of hemin transferred per second, enabling more meaningful comparisons of the transfer kinetics of the receptor mutants.

## RESULTS

### Inter-domain motions enable the LN3 unit to transiently engage Hb’s heme pocket

NMR studies of IsdH^N2N3^ in its Hb-free state (hereafter called apo-IsdH^N2N3^) revealed the presence of significant inter-domain motions between its N2 and linker domains, and suggested that these motions persist even after the receptor binds to Hb [10, 29]. To probe these domain rearrangements, we performed molecular dynamics (MD) simulations of apo-IsdH^N2N3^ and the IsdH^N2N3^:Hb complex. Three 500 ns simulations of apo-IsdH^N2N3^ were performed, revealing large scale inter-domain motions between the N2 domain and the LN3 unit that are enabled by coordinate rearrangements in the ten amino acid tether sequence (residues P466-Y475) (**Fig. S1**). Two metrics were used to quantify the inter-domain motions: the inter-domain separation between the centers of mass (COM) of the N2 and N3 domains, and the degree of twisting defined by the dihedral angle between the N2 COM, the intervening linear linker domain, and the N3 COM (**Fig. 2A**). Each 500 ns simulation showed distinct inter-domain trajectories despite starting from identical initial conformations in which the domains are separated by ∼49 Å (**Fig. 2B**). To sample longer time scale receptor motions, 20 rounds of ten x 100 ns MD simulations were performed using the FAST protocol (see methods), and trajectories were analyzed by constructing a Markov model [37]. The model provides a population estimate for 200 microstates, each corresponding to a distinct inter-domain arrangement between the N2 and LN3 units. These parameters were then used to generate a free energy landscape of inter-domain motions. In the apo-receptor the landscape is relatively flat, with most conformers accessible at room temperature; their free energies are only 2-4 kcal/mol above the ground state (**Fig. 2C**). However, a closed state in which N2 and N3 domains are proximal to one another is slightly preferred. This agrees with our original three 500 ns trajectories, which revealed that the compact state can be stable for hundreds of nanoseconds. In the apo-state the receptor also transiently samples the conformation of the Hb-bound form visualized by crystallography (star), although it represents only a small fraction of the conformers sampled in equilibrium. The MD-derived energy landscape is also compatible with an NMR model of the apo-receptor, which samples a narrower distribution of receptor conformers (indicated by black dots on the landscape) [10].

**Figure 2.**
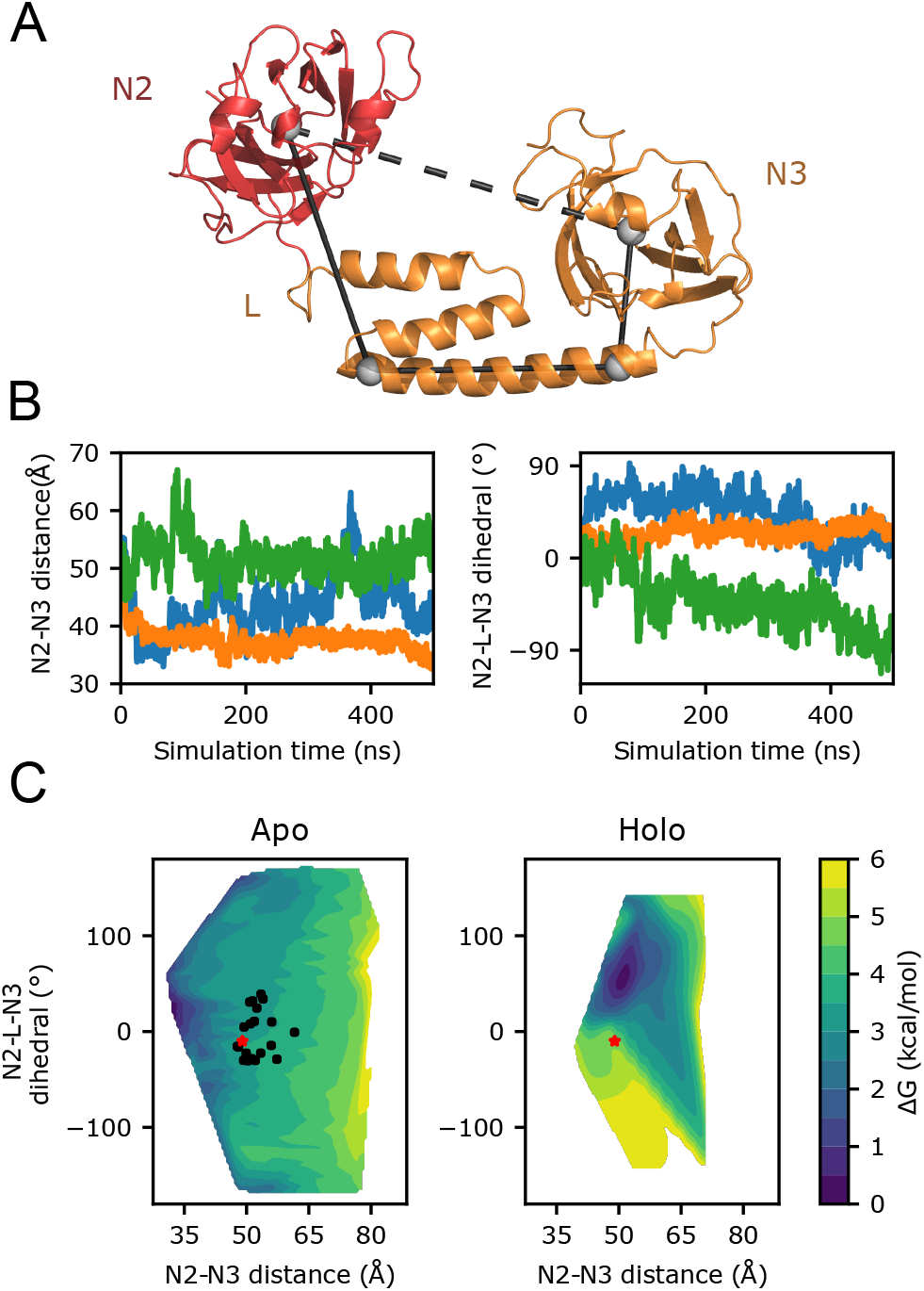
MD simulations of apo-IsdH^N2N3^ and the IsdH^N2N3^:Hb complex. (A) Visualization of the metrics used to describe interdomain motions in the receptor: (i) the distance between the centers of mass (COM) of the N2 and N3 domains (dashed line), and (ii) the dihedral angle between the COM of the domains. The angle is defined by the N2 COM, the COM of the first four and last four residues of the third helix in the linker, and the N3 COM (solid lines). (B) Time evolution plots showing inter-domain motions in three separate 500 ns simulations. Flexibility was analyzed by determining changes in the domain-domain separation (left) and twist angle (right) as described in panel A. (C) Free energy landscape parameterized by N2-N3 domain separation and twist angle for both the apo-receptor (left) and the IsdH^N2N3^:Hb complex (right). Values for these parameters were estimated from a Markov model built on multiple trajectories. In apo-IsdH^N2N3^ a wide range of inter-domain conformations are observed that sample receptor conformers that were seen in the crystal structure of the IsdH^N2N3^:Hb complex (red star) and structures of apo-IsdH^N2N3^ determined by NMR (black dots). Significant inter-domain motions persist in simulations of the IsdH^N2N3^:Hb complex, but are more restricted than those observed for apo-IsdH^N2N3^.

To explore the effects Hb binding on receptor inter-domain motions we performed MD simulations of the IsdH^N2N3^:Hb complex and constructed a Markov model. Initially, we obtained diverse apo-state configurations by coarse graining the apo-state Markov model into 75 macrostates using BACE [40], and then we selected a representative structure at random from each macrostate. A Hb dimer was then introduced into the system by aligning the N2 domain in each representative structure to the N2 domain located in the crystal structure of the IsdH^N2N3^:Hb complex in which Hb is dimeric (PDB: 4XS0). This procedure was used because the affinity of N2 for Hb’s A-helix is much stronger than the LN3 unit that contacts the heme pocket [11]. After alignment, some of the structures exhibited atomic overlap between the coordinates of the Hb dimer and the LN3 unit and were eliminated. This left approximately 50 starting structures of the IsdH^N2N3^:Hb complex that were each simulated for 100 ns, followed by further sampling using the FAST protocol and steered-MD, leading to 4.5 µs of total sampling (see methods).

An analysis of the Markov model of the MD trajectories reveals that the tether undergoes wide ranging coordinate rearrangements that enable the LN3 unit to move relative to the N2 domain and Hb. A comparison of the free energy landscapes of these inter-domain motions in the free- and Hb bound forms of receptor reveals substantial differences (**Fig. 2C**). Not only has the most stable conformation of the receptor changed, but the energy landscape is less flat and more well shaped when the receptor binds to Hb. Thus, IsdH remains flexible when in complex with Hb as predicted from NMR measurements, albeit less flexible than when it is in its apostate. Interestingly, in the MD simulations the LN3 unit receptor only transiently samples conformations that resemble the crystal structure of the extraction complex (red star), which are approximately 4 kcal/mol higher in energy than the most stable conformer. This is in marked contrast to the crystal structure complex in which the LN3 is engaged with Hb (**Fig. 1A**). A comparison of the conformer ensembles of apo-IsdH^N2N3^ and IsdH^N2N3^:Hb suggests that Hb binding occurs via a conformational selection mechanism as only a subset of apo-IsdH conformers are capable of engaging Hb via the their N2 domains without causing unfavorable atomic overlap with the remainder of the receptor. Collectively, the MD simulations reveal that the tether sequence connecting the N2 and LN3 units plays an instrumental role in orchestrating inter-domain motions in the receptor that persist even after it has bound to Hb.

### Altering the receptor’s inter-domain tether slows the rate of hemin extraction from Hb

To gain insight into the functional consequences of inter-domain motions, we measured the hemin extraction kinetics of a series of receptor variants in which the tether sequence connecting the N2 and LN3 unit was altered (**Fig. 1B**). As described previously, kinetic measurements employed a tetramer stabilized form of Hb (Hb0.1) and an IsdH^N2N3^ variant that preferentially binds to the α-globin within Hb0.1 (^α^IsdH^N2N3^, IsdH^N2N3^ containing F365Y rapid and slow changes in and A369F substitutions within its N2 domain) [12]. Using these protein reagents enables measurement of the rate of hemin extraction from only the α-subunit of tetrameric Hb, avoiding complications associated with Hb dissociation into its dimeric and monomeric forms. Hemin extraction rates were determined by tracking the change in Soret absorbance at 405 nm using a stopped-flow UV/Visible spectrophotometer (**Fig. 3A**). Hemin transfer from Hb to ^α^IsdH^N2N3^ is biphasic, with rapid and slow changes in A_405_ characterized by k_fast_ and k_slow_ rate constants, respectively (**Fig. 3B**). For fully functional receptors k_fast_ defines the rate constant describing active hemin extraction from Hb’s α-globin, whereas the value of k_slow_ characterizes transfer to the receptor that occurs via an indirect process in which Hb first releases hemin into the solvent followed by hemin binding to the apo-receptor [11]. To assess the effects of tether mutations on hemin transfer we compared the initial rate of hemin transfer (µM s^-1^), since for several of the receptor variants the transfer reaction was not completed during the time frame of the stopped-flow experiment. The kinetics data is reported in **Table S1** and was obtained as described in the Methods section.

**Figure 3.**
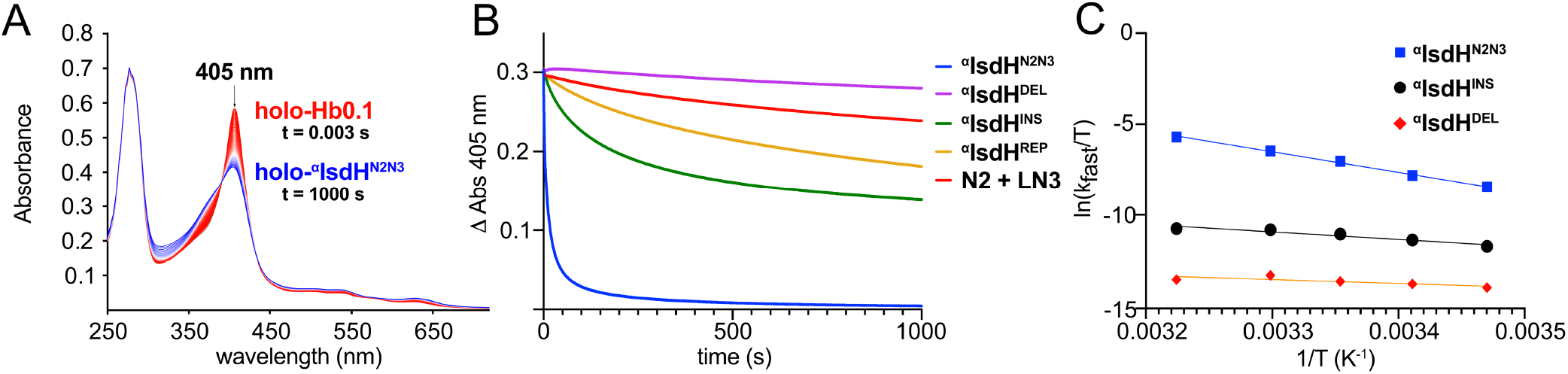
Stopped-flow measurements of hemin transfer. (A) Representative stopped-flow UV/Vis absorbance changes during the course of reaction. In the experiment apo-IsdHN2N3 (150 *µ*M) is rapidly mixed with holo-Hb0.1 (20 *µ*M) in its ferric state at 25*°*C. Traces are colored on a scale from t = 0 s (red) to t = 1000 s (blue). (B) Graph showing the change in absorbance at 405 nm after mixing holo-Hb0.1 with different tether variants of apo-αIsdH^N2N3^. Transfer measurements were performed at 25*°*C using the following receptor variants: apo-αIsdH^N2N3^, αIsdH^DEL^, αIsdH^INS^, αIsdH^REP^ and N2 + LN3 added in trans. The data shows that altering the tether significantly slows transfer, which remains incomplete even after 1000 s. (C) Eyring plots for IsdH tether variants that were used to determine the thermodynamic activation parameters (ΔH^╪^ and ΔS^╪^) of hemin transfer. The variants show distinct thermodynamic signatures indicating that they scavenge hemin via both direct and indirect mechanisms.

Rapid mixing experiments using native ^α^IsdH^N2N3^ reveal that it actively extracts hemin from Hb at a rate of 0.93 ± 0.05 µM s^-1^. This is significantly faster than the rate at which Hb0.1 spontaneously releases hemin into the solvent, consistent with an active extraction process. Housing the N2 and LN3 units within the same polypeptide is critical, since a polypeptide containing only the isolated LN3 unit (IsdH^LN3^) captures hemin from Hb0.1 435-fold slower than the native ^α^IsdH^N2N3^ receptor (0.002 ± 0.004 µM s^-1^). Adding a polypeptide encoding the N2 domain to this reaction (^α^IsdH^N2^) only marginally affects the rate of hemin capture by IsdH^LN3^, indicating that the main function of N2 is to tether the LN3 unit to Hb0.1 and its N2 binding to the A-helix does not promote hemin release.

Three receptor variants in which the tether was altered were studied: (i) ^α^IsdH^INS^, an insertion variant containing an additional ten residue (GS)_5_ segment in the tether that is located after residue P466, (ii) ^α^IsdH^DEL^, a tether deletion variant that removes residues P466-Y475, and (iii) ^α^IsdH^REP^, a variant in which amino acid sequence P466-Y475 in the tether is replaced with (GS)_5_ (**Fig. 1B**). Stopped-flow experiments reveal that replacing the tether’s amino acid sequence with (GS)_5_ or lengthening it slows hemin capture dramatically, as the ^α^IsdH^REP^ and ^α^IsdH^INS^ variants are slowed 320- and 73-fold relative to ^α^IsdH^N2N3^, respectively. ^α^IsdH^DEL^ in which the tether sequence is removed entirely is extremely slow at capturing hemin, with transfer rates 4140-fold slower than the native receptor. In fact, the rate of hemin transfer by ^α^IsdH^DEL^ is substantially slower than the intrinsic rate of spontaneous hemin loss from Hb0.1. This result suggests that ^α^IsdH^DEL^ acts to inhibit hemin loss from Hb0.1, perhaps by blocking hemin egress from Hb through steric occlusion by the misbound LN3 unit. This massive loss in activity is not caused by disruption of the tertiary structure of the receptor, as ^15^N-labeled ^α^IsdH^DEL^ and ^α^IsdH^N2N3^ exhibit similar NMR spectra. Collectively, our results show that the tether sequence serves as the nexus for inter-domain motions in the receptor and plays a critical role in coupling the destabilization of Hb’s hemin pockets with the direct extraction and capture of hemin by the N3 domain.

### The tether promotes directed inter-domain motions that orient the LN3 unit for hemin extraction

To gain in-sight into why altering the tether affects the rate of hemin extraction, we simulated the dynamics of the ^α^IsdH^INS^ and ^α^IsdH^DEL^ receptors when bound to Hb and produced a holo-state Markov model as described above for native ^α^IsdH^N2N3^. Altering the tether results in profound changes in the energetic landscapes describing inter-domain motions between the N2 and LN3 units (compare **Figs. S2** and **S3**). These differences are highlighted in **Fig. 4** which shows a probability-weighted atomic density map defining the positioning of the N3 domain during the course of each simulation. In the native ^α^IsdH^N2N3^ receptor the N3 domain in the LN3 unit primarily stays affixed to Hb’s pocket, sampling a range of conformations near Hb that resemble its positioning in the crystal structure (**Fig. 4A**, high atomic density colored red and the location of N3 in the crystal structure of the receptor complex indicated by a dashed line). Conformers in which the N3 domain is positioned distal to Hb are also sampled, but occur less frequently. In contrast, a much larger distribution of N3 positions is observed in simulations of ^α^IsdH^INS^ in which the tether sequence is lengthened, as evidenced by an expanded density cloud (**Fig. 4B**). Moreover, the most prevalent position sampled by N3 in the Hb: ^α^IsdH^INS^ complex no longer resembles the crystal structure, suggesting the tether sequence is required to properly position N3 near Hb, and explaining why its alteration slows the rate of hemin removal. Simulations of the Hb:IsdH^DEL^ complex in which the tether is removed reveal that the LN3 unit remains dynamic. Nevertheless, its positioning is biased toward a closed state in which the N3 domain is near Hb’s hemin, but translated as compared to the native receptor (**Fig. 4C**). Transfer measurements indicate that IsdH^DEL^ captures hemin very slowly, slower than the rate at which Hb spontaneously releases hemin into the solvent. This is compatible with the simulation data as the close proximity of the N3 domain relative to Hb may act to block hemin release, and because the domain is displaced, the LN3 unit may be unable to distort Hb’s F-helix to trigger hemin release.

**Figure 4.**
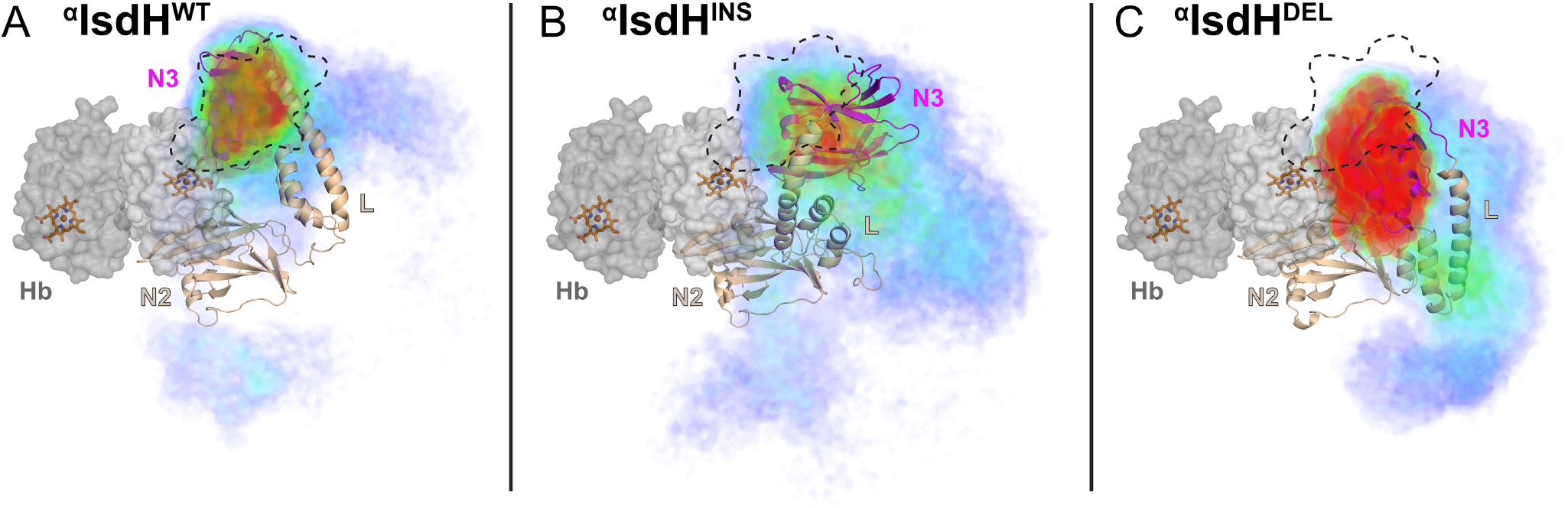
Atomic density plots describing the positioning of the N3 domain in wild-type and mutant receptors. The positioning of the N3 domain in simulations of complexes containing Hb bound to ^α^IsdH^WT^ (left), ^α^IsdH^INS^ (middle) and ^α^IsdH^DEL^ (right) are shown. A space filling model of dimeric α/β Hb is shown (gray) with its hemin molecules drawn in stick format. In each image the structure of the most populated form of the receptor is shown as a cartoon. The positioning of the N3 domain sampled in each simulation trajectory is represented as an atomic density map (blue = low density, red = high density). The positioning of the N3 domain observed in the crystal structure of the complex (PDB: 4XS0) is indicated by dashed outline (black) and presumably represents the active form of the receptor. The N3 domain in simulations containing ^α^IsdH^WT^ have the greatest degree of density overlap with the crystal structure, whereas in ^α^IsdH^INS^ the N3 density is more disperse. In ^α^IsdH^DEL^ simulations the N3 domain is translated relative to its positioning observed in the crystal structure of the receptor–Hb complex.

A Markov model of the simulation data was constructed to further probe how the tether sequence impacts the ability of the LN3 extraction unit to engage Hb’s heme pocket. The wild-type and mutant receptor-Hb trajectories were coarse grained, grouped into ∼150 microstates and a Markov model constructed that contained 25 macrostates. These macrostates were then placed into one of two categories: an associated group where the average minimum distance between Hb and the N3 domain was below 4 Å, and a dissociated group that was separated by greater than 4 Å. This definition allowed us to estimate the timescale of the N3-Hb interaction in a general manner, as the association group involves both direct transfer competent and incompetent Hb-receptor orientations. The average association transition times required for the N3 domain in each type of receptor to approach Hb’s heme pocket are shown in **Fig 5A**, providing insight into the frequency of engagement. Consistent with density plot analysis that shows that the N3 domains in the wild-type and tether deletion mutant tend to hover near Hb, in the ^α^IsdH^DEL^ and ^α^IsdH^N2N3^ proteins N3 associations occur within 7 µs.

**Figure 5.**
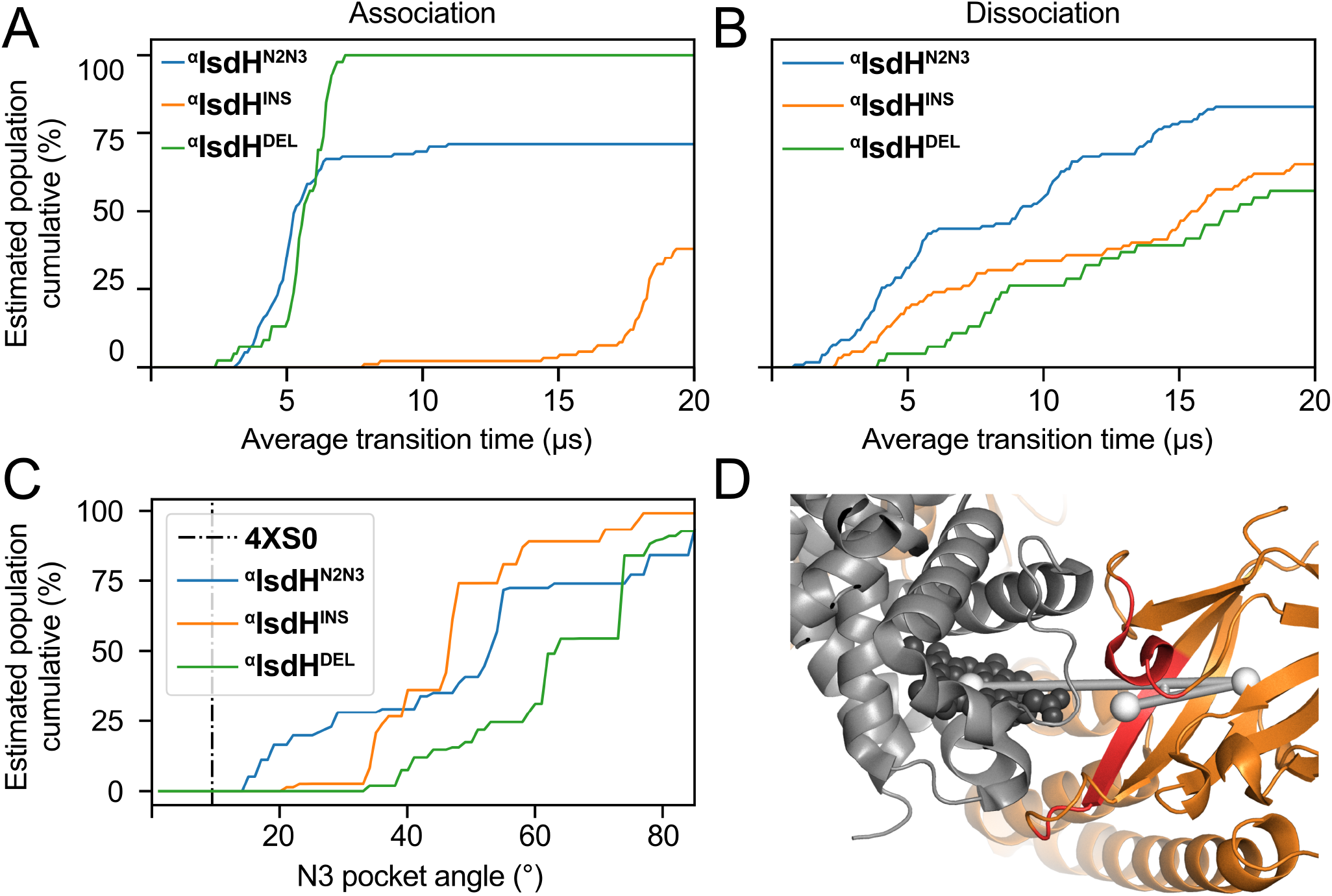
Markov model analysis of receptor proximity to Hb. Panels (A) and (B) show the time scales of association and dissociation, respectively. Values were estimated by calculating the mean first passage time between associated and dissociated macrostates. Note these charts are cumulative, meaning the percentage given is the number of transitions that occur within the timescale; for instance, 70% of association events occur within 5 µs in the case of ^α^IsdH^N2N3^. The curves are colored as follows: ^α^IsdH^WT^ (blue), ^α^IsdH^INS^ (orange) and ^α^IsdH^DEL^ (green). Panel (C) characterizes the orientation of the N3 domain relative to the Hb in the macrostates. The angle between the N3 domain and metal in the Hb bound hemin molecule was determined for each macrostate and running total calculate (left). Curves are labeled as in panels A and B. The angle was calculated from three points: hemin Fe^3+^, the center of mass of N3, and Y642 (right). As compared to the mutants the wild-type ^α^IsdH^WT^ receptor adopts orientations that more closely resemble the 2.6 Å crystal structure of the IsdH^N2N3^-Hb complex in which Hb’s heme pocket is distorted (vertical dashed line).

In the case of ^α^IsdH^DEL^ these associated states presumably represent inactive configurations in which the N3 domain is translated relative the Hb’s hemin (**Fig. 5B**). Moreover, compatible with density plots that show that the N3 domain in IsdH^INS^ samples a wider range of configurations that are removed from Hb, lengthening the tether slows the rate of N3 association with Hb, which requires on average more than 15 µs. Less substantial differences between the receptor types are observed when the average N3 dissociation times are compared, however dissociation events for both mutations occur on a longer timescale.

If the tether acts to bias inter-domain motions toward Hb, does it also impart a directional prejudice that increases the likelihood that the LN3 unit encounters Hb in an extraction competent orientation? Previous studies identified two subsites in the LN3 unit that promote hemin release from Hb by contacting its F-helix, the “L” and “N3” sub-sites [29] (**Fig. 6A**). We reasoned that LN3 conformers in the MD trajectory that were oriented similarly to the crystal structure of the IsdH^N2N3^:Hb complex would be poised to extract hemin and perhaps enriched in simulations of the wild-type receptor. To investigate this issue, we specified the orientation of receptors in the MD trajectories using three vectors whose relative positioning provides insight into the separation and orientation of the LN3 unit relative to Hb (**Fig. 6A**, purple). These included a vector connecting points within N2 domain (N2 vector), a vector through the linker domain connecting N2 to N3 (L vector) and a vector connecting points within N3 (the N3 vector). N2 and Hb are tightly bound and move with respect to one another during the simulation. Therefore, the angle between the L and N2 vectors (N2-L angle) is sensitive to LN3 motions that alter its displacement from Hb, while the dihedral angle generated by rotating about the L vector (L dihedral angle) reports on the orientation of the sub-sites within LN3 unit. **Fig. 6B** shows a plot of these angular parameters generated using the 25 macrostates that define the ensemble of receptor conformations present in the MD simulations. The crystal structure presumably represents a hemin extraction-competent configuration and has an N2-L angle and L dihedral angle of 79.2° and −60.2°, respectively (colored red in **Fig. 6B**). Each type of receptor samples a range of conformations that differ from the crystal structure based on their angle and dihedral metrics. However, inspection of the ^α^IsdH^WT^ macrostates reveals that they are enriched for conformers that resemble the crystal structure (left). The active configuration orientation for the native receptor is also evident when the vectors for each protein are visually displayed after aligning their N2 vectors (**Fig. 6B**, top). Consistent with the atomic density plots, ^α^IsdH^WT^ samples a range of conformations, but has a preference for closed configurations that resemble the crystal structure in both its distance separation and orientation of the sub-sites relative to Hb. In contrast, in the ^α^IsdH^INS^ simulations the extraction unit samples a much wider range of orientations that differ in their separation (N2-L angle) and orientation (L-dihedral angle) relative to Hb. Moreover, even though ^α^IsdH^DEL^ tends to position its LN3 unit near Hb, a large scatter in the dihedral angle indicates that its LN3 unit is misaligned with Hb. The conclusion that the tether plays a critical role in enabling the extraction unit to transiently engage Hb’s F-helix in functionally active orientations is further supported by measurements of the N3 pocket angle, a metric that measures the alignment of the heme binding pockets in Hb and the receptor (**Fig. 5C**). This analysis reveals that in simulations of the wild-type receptor ∼20% of the conformers in the trajectory have pocket angles that are within 10° of the crystal structure, whereas the insertion and deletion tether mutations rarely sample this extraction competent configuration. Thus, we conclude that the tether sequence acts to direct motions in the LN3 unit such that the unit samples conformers that are properly oriented and positioned for hemin extraction.

**Figure 6.**
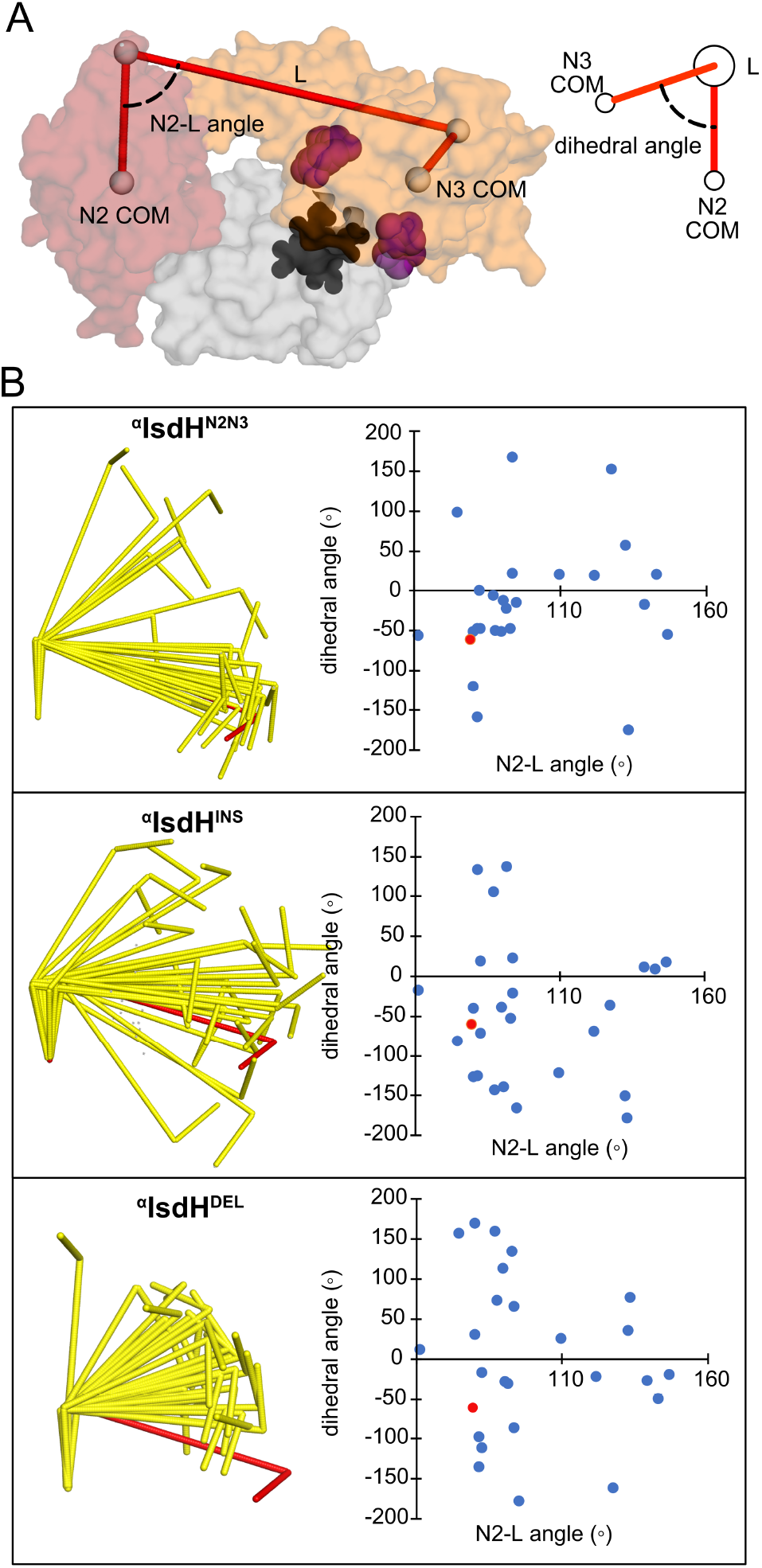
Markov model dihedral angle analysis of the receptor’s orientation relative to Hb. (A) Image showing the angular metrics that were used to represent receptor orientation. A space filling model of the IsdH^N2N3^:Hb complex is shown (PDB code: 4XS0). Color code: Hb, gray; heme, black; N2, red; LN3 unit, orange. The sub-sites are colored purple and labeled. Solid lines show the three vectors that were to calculate the dihedral angle and N2-L angle parameters. (B) Graphs comparing the dihedral orientations of native and mutant receptors in the MD simulations. (top panel) The orientations of the vectors defined in panel (A) are shown for the 25 macrostates (yellow) obtained from simulations of ^α^IsdH^N2N3^ (top), ^α^IsdH^INS^ (middle) and ^α^IsdH^DEL^ (bottom). Scatter plot showing the dihedral angle and N2-L angle metrics that report on LN3 orientation relative to Hb. For both panels shown in red is the data obtained from the crystal structure of the IsdH^N2N3^:Hb complex in which the F-helix is distorted. The data show that when the wild-type tether is present the LN3 domain has a greater tendency to sample orientations in which the “L” and “N3” sub-sites on the receptor are properly aligned with Hb’s F-helix.

### Perturbing the receptor’s inter-domain dynamics causes it to passively acquire hemin from the solvent

If altering inter-domain dynamics slows hemin capture, does it also affect the mechanism through which hemin is transferred from Hb to the receptor? To investigate this issue the temperature dependence of the rate of hemin capture for wild-type and mutant forms of the receptor were measured and interpreted using an Eyring analysis (**Table S2**). A similar analysis was performed for the hemin transfer reaction from Hb to apo-Mb, which is known to occur through an in-direct process in which Hb first releases hemin into the solvent before it is bound by apo-Mb. Based on their measured extraction velocities, the tether variants can be ranked from fastest to slowest as: ^α^IsdH^N2N3^ > ^α^IsdH^INS^ > ^α^IsdH^REP^ > ^α^IsdH^DEL^ (**Fig. 3B** and **Table S1**). Interestingly, as the receptor hemin capture efficiencies decrease ΔS^╪^ becomes progressively less favorable, trending from a favorable 14.9 cal/mol*K for ^α^IsdH^N2N3^ to an unfavorable −59.5 cal/mol*K for ^α^IsdH^DEL^ (**Fig. 7**). An opposite trend in ΔH^╪^ occurs, which becomes more favorable as the receptor become less able to rapidly capture hemin; 22.6 ± 0.7 kcal/mol (^α^IsdH^N2N3^) to 4.3 ± 1.7 kcal/mol (^α^IsdH^DEL^). The energetic profile of ^α^IsdH^DEL^ is like that observed for apo-Mb, suggesting that it is unable to actively distort Hb’s heme pocket, in-stead scavenging hemin that is released from Hb into the solvent.

**Figure 7.**
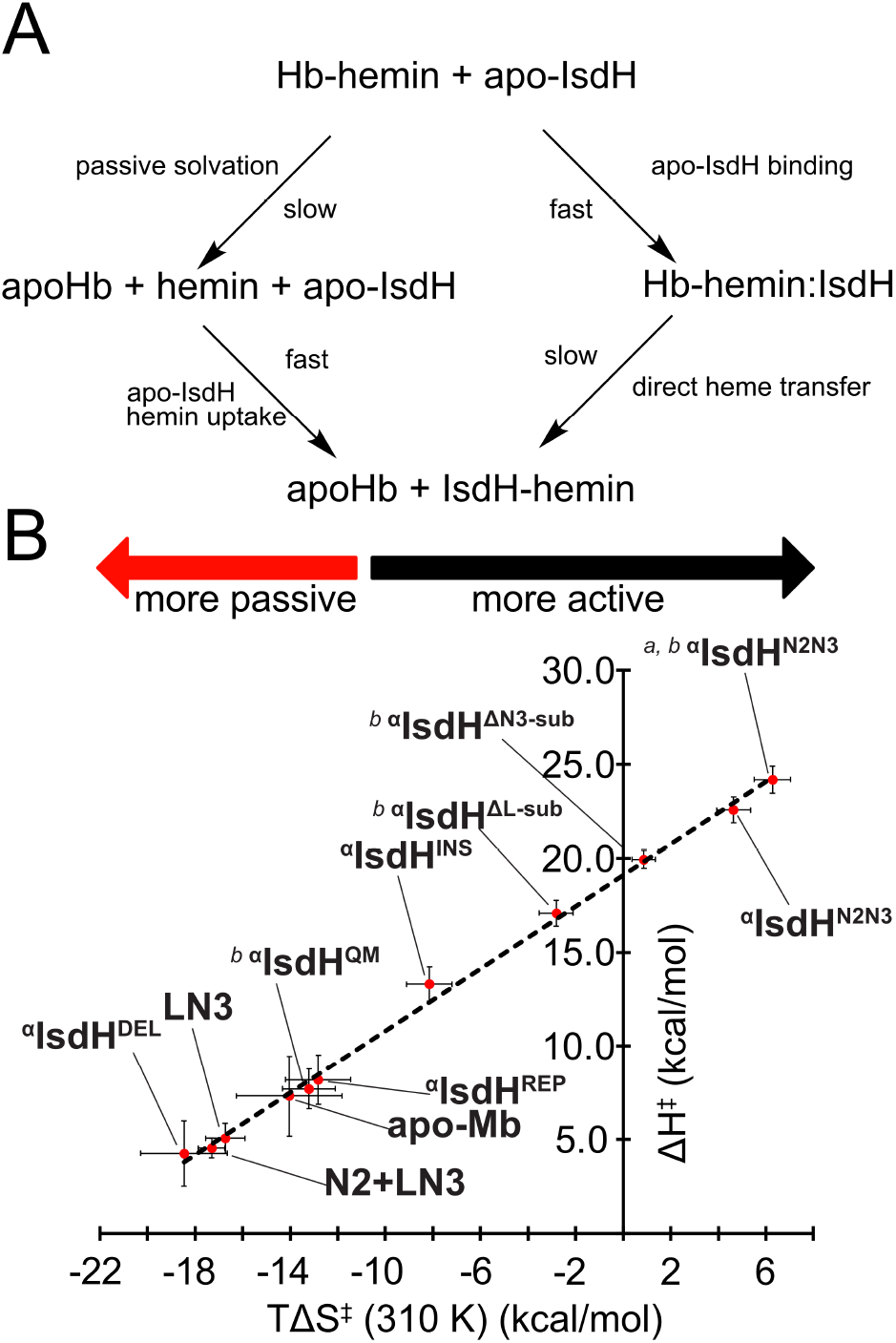
Enthalpy and entropy activation values for IsdH receptor variants. (A) Reaction schematic showing passive (left) and active (right) hemin extraction pathways from Hb. (B) Plotted values for entropy vs. enthalpy of a linear relationship between the energetic parameters. Slower mutants are characterized by unfavorable entropy of activation and relatively favorable enthalpy of activation, whereas faster mutants show more favorable entropy of activation and greater enthalpy of activation. We postulate that this relationship stems from differing proportions of direct versus passive transfer as direct hemin transfer activity is impaired. ^a^Reported in reference 11. ^b^Reported in reference 9.

## DISCUSSION

Inter-domain motions play a key role in modulating protein function by altering their affinity and specificity for ligands, as well as governing the efficiencies with which they catalyze chemical transformations [47, 48]. Domains are frequently connected by short polypeptide tethers, whose length and amino acid composition constrains their range of motion [49, 50]. Here we show using MD simulations and quantitative measurements of hemin transfer that a flexible inter-domain tether within the IsdH receptor plays a critical role in controlling domain motions that are needed to extract hemin from Hb. Simulations of the native complex reveal that it is dynamic, with coordinate rearrangements in the tether enabling the LN3 unit to adopt a range of conformations relative to Hb, while the N2 domain remains affixed to the globin A-helix. Interestingly, these inter-domain motions are biased, such that within the cloud of receptor conformers there is an enrichment for LN3 units that are positioned near Hb’s heme pocket (**Figs. 2, 4**) that are also properly oriented to distort its F-helix (**Figs. 5, 6**). The specific amino acid sequence of the tether is important for function, as a ^α^IsdH^REP^ receptor variant in which the tether is replaced with glycine and serine residues acquires heme 320-fold slower than the wild-type receptor. The finding that tether residues enable the low-affinity LN3 unit to move even after the receptor binds to Hb is consistent with NMR studies [10]. It is also compatible with the crystal structure of IsdH^N2N3^-Hb, as electron density for 7 of the 10 residues in the tether (P466-E472) were missing suggesting that they may be flexible [26, 12]. However, in the crystal structure of the complex, the LN3 adopts a single conformation in which it is affixed to Hb, presumably because crystal packing interactions restrict its mobility.

Our results suggest that within the dynamic complex the LN3 unit repeatedly engages Hb, but it only infrequently forms extraction competent configurations in which heme is removed. Comparison of the extraction kinetics of the native receptor and an isolated LN3 unit reveals that the motional bias imparted by the native tether increases the effective concentration of the LN3 unit near Hb’s heme to ∼15 mM. This presumably drives the formation of the weaker LN3-Hb interface in which Hb’s heme pocket is distorted. A Markov model of the simulation data reveals that the LN3 unit samples a range of configurations that generally resemble the crystal structure. Using the LN3 position in the crystal structure in which sub-sites are aligned with Hb to define “transfer competence”, the model predicts that only 5% of complexes in equilibrium are competent for direct transfer, which is in agreement with previous NMR results [29]. Based on the mean first passage times of LN3, Hb association and dissociation events occur within ∼10 µs (**Fig. 5**). Using the rate constant of direct hemin transfer (*k*_fast_) for ^α^IsdH^N2N3^, the mean lifetime (τ) of the functional ^α^IsdH^N2N3^-Hb complex under pseudo-first order conditions is ∼1 s. In comparing these values, we postulate that LN3 can potentially undergo ∼10^5^ association and dissociation events prior to successfully extracting hemin. This implies that only a small subset of LN3:Hb association events within the dynamic complex satisfy conditions needed to successfully transfer heme from Hb to the receptor.

The tether connecting the N2 and LN3 units directs inter-domain motions that are important for heme extraction. This is evident from studies of ^α^IsdH^INS^, a receptor variant that contains a (GS)_5_ insertion after residue P466 in the tether. Relative to the wild-type protein it exhibits a large 73-fold reduction in the transfer rate, which coincides with the results of MD simulations that reveal that the dynamic LN3 unit samples a much larger, more isoenergetic conformational landscape in which the orientational constraints imposed by the native tether sequence are lost (**Figs. 4-6**). The increased motional freedom caused by the insertion also extends the mean first passage time needed to associate with Hb’s heme pocket from 7 to >14 µs, resulting in greater exploration of LN3 disassociated states that are unable to extract heme (**Fig. 5**). In ^α^IsdH^DEL^ the tether is deleted entirely. This rather aggressive mutation still yields a well-folded protein as determined by ^15^N-HSQC experiments, but causes a very substantial ∼4100-fold reduction in the rate of hemin capture. In the ^α^IsdH^DEL^ the receptor’s ability to orient its LN3 unit for productive Hb engagement is lost, as both the positioning of the LN3 unit (**Fig. 3**) and orientation relative to Hb (**Fig. 5**) are altered as compared to the native receptor.

An analysis of the activation energies of hemin transfer reveals that altering the tether causes the receptor to capture hemin from Hb via an indirect process in which hemin is first released into the solvent before being captured by the receptor (**Fig. 7A**). For a series of receptor variants, a plot of their activation energies reveals that slower transfer rates relate to ΔS^╪^ becoming more negative and ΔH^╪^ becoming smaller in magnitude. Both parameters are correlated with the degree of receptor impairment and the correlation holds true for mutant receptors with either altered tethers (^α^IsdH^REP, α^IsdH^DEL^, and ^α^IsdH^INS^) or amino acid substitutions in receptor sub-sites that directly contact Hb’s F-helix in the crystal structure (^α^IsdH^ΔL-sub^, ^α^IsdH^ΔN3-sub^ and ^α^IsdH^QM^) (**Fig. 7B**). Transfer of hemin from Hb to apo-Mb occurs through indirect mechanism and exhibits similar activation energies as the most defective IsdH receptors. Therefore, we postulate that that in both the tether and sub-site receptor mutants, an increasing proportion of hemin is transferred via an indirect mechanism wherein hemin is first released from Hb into the solvent. Presumably in the indirect process hemin solvation causes the large unfavorable change in ΔS^╪^, whereas the direct hemin transfer reaction is enthalpically limited because the receptor must distort Hb’s F-helix. Thus, combined with the MD results, the transfer data suggest that directed inter-domain motions orchestrated by the tether sequence are needed to populate conformers that provide a route for direct hemin transfer between Hb and N3.

A comparison of the tether conformations sampled during the simulations provides insight into how it directs restricted, and functionally important LN3 motions. The primary sequence of the tether (^466^PDDYVDEETY^475^) is notably populated with carboxylate sidechains that are documented to induce disorder [51]. In simulations of the native IsdH-Hb complex these sidechains frequently participate in intra-tether hydrogen bonding interactions with the polypeptide backbone that bias the tether’s conformation toward a compact state. Interestingly, in many of these condensed configurations the side chain of Y475 in the tether forms sustained interactions with a hydrophobic patch on the N2 domain thereby constraining the LN3 unit so that it is positioned near Hb’s heme pocket. This closed state is captured in the crystal structure of the complex where the Y475 aromatic ring is docked against a relatively hydrophobic surface on the N2 domain formed by P351 and P374, and poised to donate a hydrogen bond to E375 (**Fig. 8**). This constrained configuration is only transiently sampled during the simulations explaining why the LN3 unit exhibits directed motions (**Fig. 4**) and why the tether mutants remove heme slowly as they disrupt this interaction. The tether sequence in IsdB also contains conserved tyrosine residue (Y273 in IsdB numbering) and is enriched with anionic residues (**Fig. 1A**), suggesting that similar tether-N2 contacts will bias its LN3 unit toward conformations that are poised to extract hemin.

**Figure 8.**
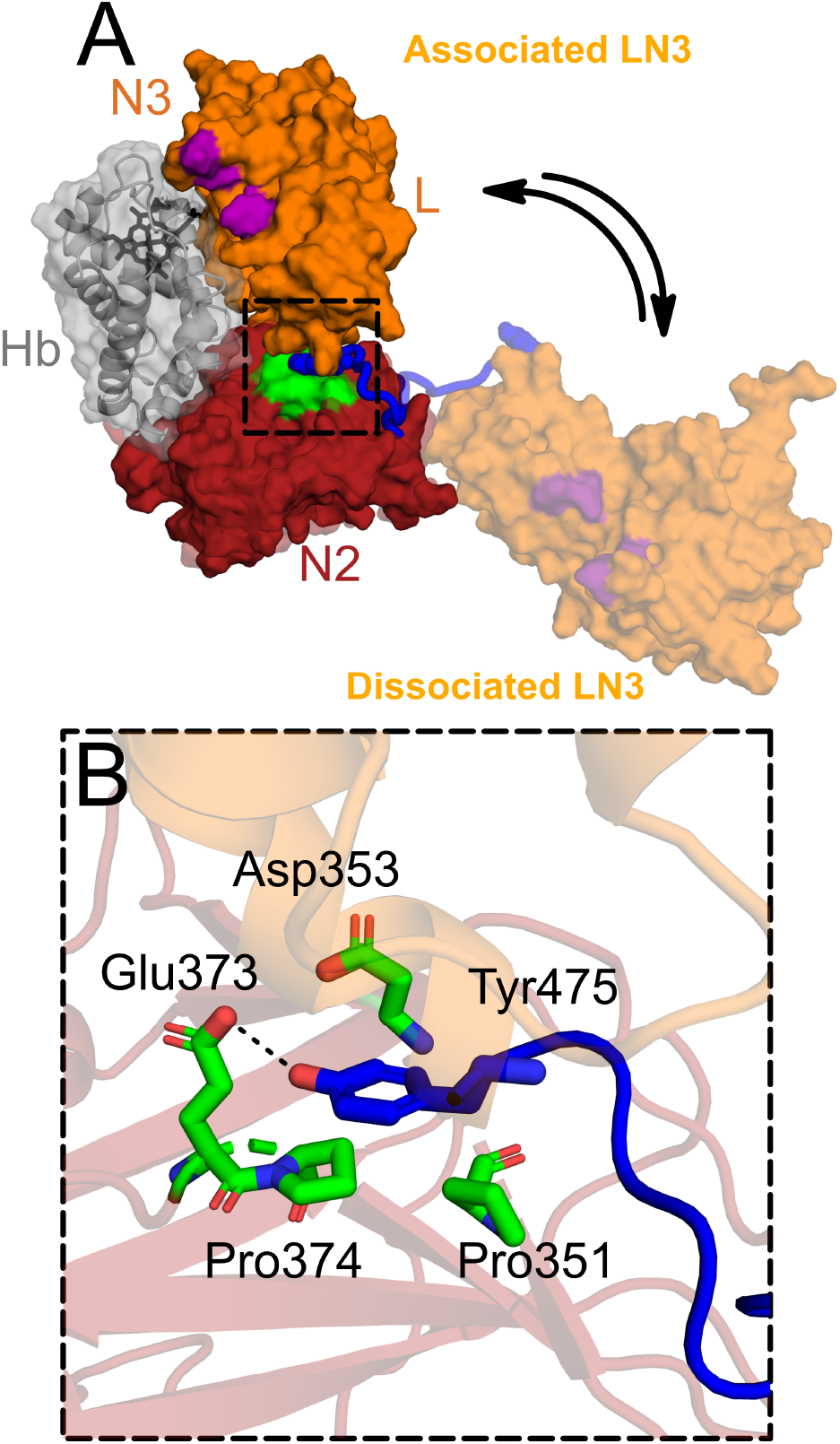
Model of on-target LN3 engagement A) Two macrostates derived from simulations of ^α^IsdH^N2N3^:Hb demonstrate the LN3 unit in disengaged (light orange) and engaged (orange) states, which reveal that Y475 (blue spheres) defines the primary contact between the tether and a conserved solvent-exposed patch on N2 (green). This interaction enforces an inter-domain geometry that positions the LN3 subsites (purple) in proximity with the heme-binding site of Hb (gray). B) A molecular model of the docked Tyr475 (blue sticks) with the cognate conserved patch residues on N2.

Domains connected by flexible tethers are often thought to exhibit disordered motions in which they freely re-orient with respect to one another. In this study we have shown that despite exhibiting a high degree of conformational flexibility, a short inter-domain tether in IsdH nevertheless restricts and directs motions of its heme extraction unit when the receptor is bound to Hb. These motions enable the LN3 extraction unit to populate a low affinity interface in which Hb’s heme pocket is transiently distorted. Altering the tether misa-ligns these motions, reducing the rate of direct hemin scavenging and causing the receptor to slowly acquire hemin through an indirect process. The functionally important motions in IsdH are akin to those observed in protein chaperones, polyketide synthases, and prolyl isomerases [49, 50, 52, 53], which also employ inter-domain tethers that determine the frequency and geometry of molecular interactions. In IsdH these motions are likely biologically important, as they enable *S. aureus* to overcome Hb’s strong affinity for its hemin molecules without the need to form a high affinity, long-lived receptor-Hb complex. In the well-studied HasA-HasR transfer complex from *Serratia marcescens*, hemin is transferred by forming a high affinity complex in which the protein-protein binding free energy is used to weaken HasA’s contacts with hemin [54–56]. In contrast, both the IsdB and IsdH receptors bind to Hb with modest affinity and harness inter-domain dynamics to facilitate heme removal. Extraction within this weak and dynamic complex presumably maximizes the rate of heme flow into the cell, enabling repeated rounds of Hb capture and heme removal to occur on the microbial surface.

## ASSOCIATED CONTENT

### Supporting Information

Supporting Information: Tables and additional figures mentioned in the text.

## AUTHOR INFORMATION

### Author Contributions

The manuscript was written through contributions of all authors. All authors have given approval to the final version of the manuscript.

### Funding Sources

Experimental work in the Clubb group was funded by the National Science Foundation MCB-1716948 and the National Institutes of Health R01A155217 and R01A116828. Work in the Wereszczynski group was funded by the National Science Foundation MCB-1716099 and the National Institutes of Health R35GM119647. This work used the Comet supercomputer at the San Diego Supercomputing Center (SDSC) through an allocation provided by the Extreme Science and Engineering Discovery Environment (XSEDE), which is supported by National Science Foundation ACI-1548562 [57].

## ACKNOWLEDGMENT

We thank members of the Clubb and Wereszczynski laboratories for useful discussions.

## ABBREVIATIONS

NEAT: NEAr iron Transporter
Hb: hemoglobin

## Insert Table of Contents artwork here

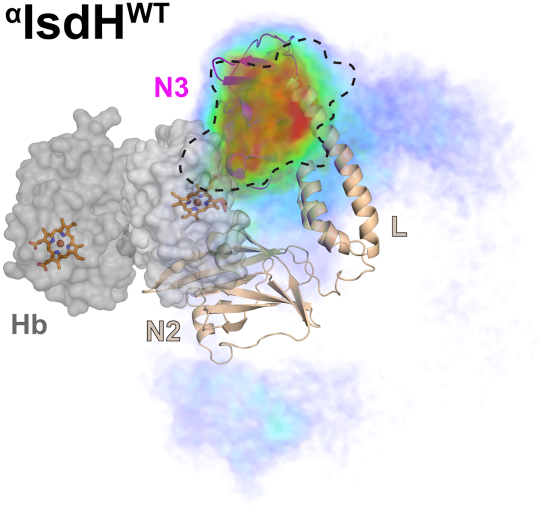

## Notes

### Competing Interest Statement

The authors have declared no competing interest.

## REFERENCES

1. Archer GL (1998) Staphylococcus aureus: A Well-Armed Pathogen. Clinical Infectious Diseases 26:1179–1181. https://doi.org/10.1086/520289

2. Klevens RM, Morrison MA, Nadle J, et al (2007) Invasive Methicillin-Resistant Staphylococcus aureus Infections in the United States. JAMA 298:1763–1771. https://doi.org/10.1001/jama.298.15.1763

3. Lee AS, de Lencastre H, Garau J, et al (2018) Methicil-lin-resistant Staphylococcus aureus. Nature Reviews Disease Pri-mers 4:. https://doi.org/10.1038/nrdp.2018.33

4. Skaar EP, Humayun M, Bae T, et al (2004) Iron-Source Preference of Staphylococcus aureus Infections. Science 305:1626–1628. https://doi.org/10.1126/science.1099930

5. Maresso AW, Schneewind O (2006) Iron Acquisition and Transport in Staphylococcus aureus. BioMetals 19:193–203. https://doi.org/10.1007/s10534-005-4863-7

6. Mazmanian SK, Ton-That H, Su K, Schneewind O (2002) An iron-regulated sortase anchors a class of surface protein during Staphylococcus aureus pathogenesis. Proceedings of the National Academy of Sciences 99:2293–2298. https://doi.org/10.1073/pnas.032523999

7. Haley KP, Skaar EP (2012) A battle for iron: host sequestration and Staphylococcus aureus acquisition. Microbes and infection 14:217–227. https://doi.org/10.1016/j.micinf.2011.11.001

8. Zhu H, Li D, Liu M, et al (2014) Non-heme-binding domains and segments of the Staphylococcus aureus IsdB protein critically contribute to the kinetics and equilibrium of heme acquisition from methemoglobin. PLoS One 9:e100744. https://doi.org/10.1371/journal.pone.0100744

9. Spirig T, Malmirchegini GR, Zhang J, et al (2013) Staphylococcus aureus uses a novel multidomain receptor to break apart human hemoglobin and steal its heme. J Biol Chem 288:1065–1078. https://doi.org/10.1074/jbc.M112.419119

10. Sjodt M, Macdonald R, Spirig T, et al (2016) The PRE-Derived NMR Model of the 38.8-kDa Tri-Domain IsdH Protein from Staphylococcus aureus Suggests That It Adaptively Recognizes Human Hemoglobin. Journal of Molecular Biology 428:1107–1129. https://doi.org/10.1016/j.jmb.2015.02.008

11. Sjodt M, Macdonald R, Marshall JD, et al (2018) Energetics underlying hemin extraction from human hemoglobin by Staphylococcus aureus. Journal of Biological Chemistry 293:6942–6957. https://doi.org/10.1074/jbc.RA117.000803

12. Dickson CF, Jacques DA, Clubb RT, et al (2015) The structure of haemoglobin bound to the haemoglobin receptor IsdH from ıt Staphylococcus aureus shows disruption of the native α-globin haem pocket. Acta Crystallographica Section D 71:1295–1306. https://doi.org/10.1107/S1399004715005817

13. Pishchany G, Dickey SE, Skaar EP (2009) Subcellular Localization of the Staphylococcus aureus Heme Iron Transport Components IsdA and IsdB. Infection and Immunity 77:2624–2634. https://doi.org/10.1128/iai.01531-08

14. Villareal VA, Spirig T, Robson SA, et al (2011) Transient Weak Protein–Protein Complexes Transfer Heme Across the Cell Wall of Staphylococcus aureus. Journal of the American Chemical Society 133:14176–14179. https://doi.org/10.1021/ja203805b

15. Zhu H, Liu M, Lei B (2008) The surface protein Shr of Streptococcus pyogenes binds heme and transfers it to the streptococcal heme-binding protein Shp. BMC microbiology 8:15–15. https://doi.org/10.1186/1471-2180-8-15

16. Reniere ML, Ukpabi GN, Harry SR, et al (2010) The IsdG-family of haem oxygenases degrades haem to a novel chromophore. Mol Microbiol 75:1529–1538. https://doi.org/10.1111/j.1365-2958.2010.07076.x

17. Matsui T, Nambu S, Ono Y, et al (2013) Heme Degradation by Staphylococcus aureus IsdG and IsdI Liberates Formaldehyde Rather Than Carbon Monoxide. Biochemistry 52:3025–3027. https://doi.org/10.1021/bi400382p

18. Andrade MA, Ciccarelli FD, Perez-Iratxeta C, Bork P (2002) NEAT: a domain duplicated in genes near the components of a putative Fe3+ siderophore transporter from Gram-positive pathogenic bacteria. Genome biology 3:RESEARCH0047– RESEARCH0047. https://doi.org/10.1186/gb-2002-3-9-re-search0047

19. Ellis-Guardiola K, Mahoney BJ, Clubb RT (2020) NEAr Transporter (NEAT) Domains: Unique Surface Displayed Heme Chaperones That Enable Gram-Positive Bacteria to Capture Heme-Iron From Hemoglobin. Front Microbiol 11:607679. https://doi.org/10.3389/fmicb.2020.607679

20. Pishchany G, Sheldon JR, Dickson CF, et al (2013) IsdB-dependent Hemoglobin Binding Is Required for Acquisition of Heme by Staphylococcus aureus. The Journal of Infectious Diseases 209:1764–1772. https://doi.org/10.1093/infdis/jit817

21. Torres VJ, Pishchany G, Humayun M, et al (2006) Staphylococcus aureus IsdB is a hemoglobin receptor required for heme iron utilization. J Bacteriol 188:8421–8429. https://doi.org/10.1128/JB.01335-06

22. Sheldon JR, Heinrichs DE (2015) Recent developments in understanding the iron acquisition strategies of gram positive pathogens. FEMS Microbiology Reviews 39:592–630. https://doi.org/10.1093/femsre/fuv009

23. Nobles CL, Maresso AW (2011) The theft of host heme by Gram-positive pathogenic bacteria. Metallomics 3:788–796. https://doi.org/10.1039/c1mt00047k

24. Pilpa RM, Robson SA, Villareal VA, et al (2009) Functionally distinct NEAT (NEAr Transporter) domains within the Staphylococcus aureus IsdH/HarA protein extract heme from methemoglobin. J Biol Chem 284:1166–1176. https://doi.org/10.1074/jbc.M806007200

25. Watanabe M, Tanaka Y, Suenaga A, et al (2008) Structural basis for multimeric heme complexation through a specific protein-heme interaction: the case of the third neat domain of IsdH from Staphylococcus aureus. J Biol Chem 283:28649–28659. https://doi.org/10.1074/jbc.M803383200

26. Dickson CF, Kumar KK, Jacques DA, et al (2014) Structure of the Hemoglobin-IsdH Complex Reveals the Molecular Basis of Iron Capture by Staphylococcus aureus. Journal of Biological Chemistry 289:6728–6738. https://doi.org/10.1074/jbc.M113.545566

27. Bowden CFM, Chan ACK, Li EJW, et al (2018) Structure–function analyses reveal key features in Staphylococcus aureus IsdB-associated unfolding of the heme-binding pocket of human hemoglobin. Journal of Biological Chemistry 293:177–190. https://doi.org/10.1074/jbc.M117.806562

28. Mikkelsen JH, Runager K, Andersen CBF (2020) The human protein haptoglobin inhibits IsdH-mediated heme-sequestering by Staphylococcus aureus. Journal of Biological Chemistry 295:1781–1791. https://doi.org/10.1074/jbc.RA119.011612

29. Ellis-Guardiola K, Joseph Clayton, Pham C, et al (2020) The Staphylococcus aureus IsdH Receptor Forms a Dynamic Complex with Human Hemoglobin that Triggers Heme Release via Two Distinct Hot Spots. Journal of Molecular Biology 432:1064–1082. https://doi.org/10.1016/j.jmb.2019.12.023

30. Joung IS, Cheatham TE (2008) Determination of Alkali and Halide Monovalent Ion Parameters for Use in Explicitly Solvated Biomolecular Simulations. J Phys Chem B 112:9020–9041. https://doi.org/10.1021/jp8001614

31. Jorgensen WL, Chandrasekhar J, Madura JD, et al (1983) Comparison of simple potential functions for simulating liquid water. The Journal of Chemical Physics 79:926–935. https://doi.org/10.1063/1.445869

32. Maier JA, Martinez C, Kasavajhala K, et al (2015) ff14SB: Improving the Accuracy of Protein Side Chain and Backbone Parameters from ff99SB. Journal of Chemical Theory and Computation 11:3696–3713. https://doi.org/10.1021/acs.jctc.5b00255

33. Loncharich RJ, Brooks BR, Pastor RW (1992) Langevin dynamics of peptides: The frictional dependence of isomerization rates ofN-acetylalanyl-N?-methylamide. Biopolymers 32:523–535. https://doi.org/10.1002/bip.360320508

34. Åqvist J, Wennerström P, Nervall M, et al (2004) Molecular dynamics simulations of water and biomolecules with a Monte Carlo constant pressure algorithm. Chemical Physics Letters 384:288–294. https://doi.org/10.1016/j.cplett.2003.12.039

35. Case DA, Ben-Shalom IY, Brozell SR, et al (2018) AMBER 2018. University of California San Francisco, San Francisco

36. Roe DR, Cheatham TE (2013) PTRAJ and CPPTRAJ: Software for Processing and Analysis of Molecular Dynamics Trajectory Data. Journal of Chemical Theory and Computation 9:3084–3095. https://doi.org/10.1021/ct400341p

37. Zimmerman MI, Bowman GR (2015) FAST Conformational Searches by Balancing Exploration/Exploitation Trade-Offs. Journal of Chemical Theory and Computation 11:5747–5757. https://doi.org/10.1021/acs.jctc.5b00737

38. Harrigan MP, Sultan MM, Hernández CX, et al (2017) MSMBuilder: Statistical Models for Biomolecular Dynamics. Biophysical Journal 112:10–15. https://doi.org/10.1016/j.bpj.2016.10.042

39. Schördinger, LLC (2015) The PyMOL Molecular Graphics System, Version 2.3

40. Bowman GR (2012) Improved coarse-graining of Markov state models via explicit consideration of statistical uncertainty. J Chem Phys 137:134111. https://doi.org/10.1063/1.4755751

41. Phillips JC, Braun R, Wang W, et al (2005) Scalable molecular dynamics with NAMD. J Comput Chem 26:1781–1802. https://doi.org/10.1002/jcc.20289

42. Fiorin G, Klein ML, Hénin J (2013) Using collective variables to drive molecular dynamics simulations. null 111:3345–3362. https://doi.org/10.1080/00268976.2013.813594

43. Kim DE, Chivian D, Baker D (2004) Protein structure prediction and analysis using the Robetta server. Nucleic Acids Res 32:W526–31. https://doi.org/10.1093/nar/gkh468

44. Hargrove MS, Singleton EW, Quillin ML, et al (1994) His64(E7)–>Tyr apomyoglobin as a reagent for measuring rates of hemin dissociation. Journal of Biological Chemistry 269:4207–4214. https://doi.org/10.1016/S0021-9258(17)41764-9

45. Pilpa RM, Fadeev EA, Villareal VA, et al (2006) Solution structure of the NEAT (NEAr Transporter) domain from IsdH/HarA: the human hemoglobin receptor in Staphylococcus aureus. J Mol Biol 360:435–447. https://doi.org/10.1016/j.jmb.2006.05.019

46. Ellis-Guardiola K, Soule J, Clubb ART (2021) Methods for the Extraction of Heme Prosthetic Groups from Hemoproteins. Bio Protoc 11:e4156. https://doi.org/10.21769/BioProtoc.4156

47. Henzler-Wildman K, Kern D (2007) Dynamic personalities of proteins. Nature 450:964–972. https://doi.org/10.1038/nature06522

48. Agarwal PK (2019) A Biophysical Perspective on Enzyme Catalysis. Biochemistry 58:438–449. https://doi.org/10.1021/acs.biochem.8b01004

49. Gokhale RS, Khosla C (2000) Role of linkers in communication between protein modules. Curr Opin Chem Biol 4:22–27. https://doi.org/10.1016/s1367-5931(99)00046-0

50. Gokhale RS, Tsuji SY, Cane DE, Khosla C (1999) Dissecting and exploiting intermodular communication in polyketide synthases. Science 284:482–485. https://doi.org/10.1126/science.284.5413.482

51. van der Lee R, Buljan M, Lang B, et al (2014) Classification of intrinsically disordered regions and proteins. Chem Rev 114:6589–6631. https://doi.org/10.1021/cr400525m

52. Bouchard JJ, Xia J, Case DA, Peng JW (2018) Enhanced Sampling of Interdomain Motion Using Map-Restrained Langevin Dynamics and NMR: Application to Pin1. Journal of Molecular Biology 430:2164–2180. https://doi.org/10.1016/j.jmb.2018.05.007

53. Zhu W, Li Y, Liu M, et al (2019) Uncorrelated Effect of Interdomain Contact on Pin1 Isomerase Activity Reveals Positive Catalytic Cooperativity. J Phys Chem Lett 10:1272–1278. https://doi.org/10.1021/acs.jpclett.9b00052

54. Izadi-Pruneyre N, Huché F, Lukat-Rodgers GS, et al (2006) The Heme Transfer from the Soluble HasA Hemophore to Its Membrane-bound Receptor HasR Is Driven by Protein-Protein Interaction from a High to a Lower Affinity Binding Site. Journal of Biological Chemistry 281:25541–25550. https://doi.org/10.1074/jbc.M603698200

55. Krieg S, Huche F, Diederichs K, et al (2009) Heme uptake across the outer membrane as revealed by crystal structures of the receptor-hemophore complex. Proceedings of the National Academy of Sciences 106:1045–1050. https://doi.org/10.1073/pnas.0809406106

56. Exner TE, Becker S, Becker S, et al (2020) Binding of HasA by its transmembrane receptor HasR follows a conformational funnel mechanism. Eur Biophys J 49:39–57. https://doi.org/10.1007/s00249-019-01411-1

57. Towns J, Cockerill T, Dahan M, et al (2014) XSEDE: Accelerating Scientific Discovery. Comput Sci Eng 16:62–74. https://doi.org/10.1109/MCSE.2014.80

