## Supporting Information for "Directed inter-domain motions enable the IsdH *Staphylococcus aureus* receptor to rapidly extract heme from human hemoglobin"

[a]Department of Physics and Center for Molecular Study of Condensed Soft Matter, Illinois Institute of Technology, Chicago, IL 60616, USA; [b]UCLA Department of Chemistry and Biochemistry, [c]UCLA-DOE Institute of Genomics and Proteomics and [d]Molecular Biology Institute. University of California, Los Angeles, 611 Charles Young Drive East, Los Angeles, CA 90095, USA

| Receptor | $k_{fast}$ | % WT span | Initial rate | Fold rate reduction | Meff | |
|---|---|---|---|---|---|---|
| WT $^{\alpha}$IsdH$^{N2N3}$ | 0.355 ± 0.001 $s^{-1}$ | 100% ± 1% | 0.929 ± 0.046 μM/s | 1 | 15.7 mM | This study |
| $^{\alpha}$IsdH$^{INS}$ | 0.012 ± 0.001 $s^{-1}$ | 56% ± 2% | 0.011 ± 0.001 μM/s | 73x | 0.54 mM | This study |
| $^{\alpha}$IsdH$^{DEL}$ | 0.001 ± 0.001 $s^{-1}$ | 8% ± 2% | 0.001 ± 0.001 μM/s | 4140x | 0.04 mM | This study |
| $^{\alpha}$IsdH$^{REP}$ | 0.003 ± 0.001 $s^{-1}$ | 41% ± 2% | 0.003 ± 0.001 μM/s | 320x | 0.13 mM | This study |
| $^{\alpha}$IsdH$^{N2}$ + IsdH$^{LN3}$ | 0.002 ± 0.001 $s^{-1}$ | 23% ± 10% | 0.001 ± 0.001 μM/s | 600x | 0.09 mM | This study |
| IsdH$^{LN3}$ | 0.003 ± 0.001$s^{-1}$ | 28% ± 3% | 0.002 ± 0.004 μM/s | 435x | 0.15 mM | This study |
| $^{a}$ $^{\alpha}$IsdH$^{\Delta sub\text{-}L}$ | 0.027 ± 0.001$s^{-1}$ | 74% ± 23% | 0.039 ± 0.013 μM/s | 21x | N/A | 1 |
| $^{a}$ $^{\alpha}$IsdH$^{\Delta sub\text{-}N3}$ | 0.065 ± 0.001 $s^{-1}$ | 102% ± 4% | 0.177 ± 0.009 μM/s | 5x | N/A | 1 |
| $^{a}$ $^{\alpha}$IsdH$^{QM}$ | 0.006 ± 0.001 $s^{-1}$ | 49% ± 6% | 0.006 ± 0.002 μM/s | 142x | N/A | 1 |
| Mb$^{H64Y/V68F}$ * | 0.004 ± 0.002 $s^{-1}$ | N/A | | | N/A | This study |

**SI Table 1.** Kinetic parameters characterizing the rate of hemin extraction from human hemoglobin by IsdH receptor variants. Kinetic data was obtained by monitoring changes in UV/Vis absorbance at 450 nm ($\Delta A_{405}$) upon rapidly mixing 5 μM Hb0.1 with 150 μM of the receptor variant (apo-form). $\Delta A_{405}$ absorbance curves were fit as described in the Methods section. Spontaneous hemin release from Hb0.1 was also measured using an apo-myoglobin variant (Mb$^{H64Y/V68F}$) by monitoring changes in absorbance at 600 nm as described previously [50]. [a]Data reported in ref. 34.

| mutant | $\Delta H^{\ddagger}$ (kcal/mol) | $T*\Delta S^{\ddagger}$ (kcal/mol) (95% CI) | $\Delta S^{\ddagger}$ (cal/mol*K) (95% CI) | $\Delta G^{\ddagger}$ (37°C) (kcal/mol) |
|---|---|---|---|---|
| WT $^{\alpha}$IsdH$^{N2N3}$ | 24.2 ± 0.7 | 6.3 ± 0.7 | 20.3 ± 2.3 | 17.9 ± 1.0 |
| WT $^{\alpha}$IsdH$^{N2N3}$ (Sjodt et al.) | 22.6 ± 0.7 | 4.6 ± 0.8 | 14.9 ± 2.5 | 17.9 ± 1.1 |
| $^{\alpha}$IsdH$^{INS}$ | 8.2 ± 1.3 | -12.8 ± 1.4 | -41.3 ± 4.4 | 21.5 ± 1.9 |
| $^{\alpha}$IsdH$^{DEL}$ | 4.3 ± 1.7 | -18.5 ± 1.8 | -59.5 ± 5.6 | 22.7 ± 2.5 |
| $^{\alpha}$IsdH$^{REP}$ | 13.3 ± 0.9 | -8.1 ± 1.0 | -26.3 ± 3.1 | 21.5 ± 1.3 |
| $^{\alpha}$IsdH$^{N2}$ + IsdH$^{LN3}$ | 4.5 ± 0.5 | -17.3 ± 0.6 | -55.8 ± 1.8 | 21.8 ± 0.8 |
| IsdH$^{LN3}$ | 5.1 ± 0.8 | -16.7 ± 0.8 | -53.9 ± 2.7 | 21.8 ± 1.2 |
| $^{\alpha}$IsdH$^{\Delta sub-L}$ | 17.1 ± 0.7 | -2.8 ± 0.7 | -9.1 ± 2.3 | 19.9 ± 1.0 |
| $^{\alpha}$IsdH$^{\Delta sub-N3}$ | 19.9 ± 0.5 | 0.9 ± 0.5 | 2.8 ± 1.6 | 19.1 ± 0.7 |
| $^{\alpha}$IsdH$^{QM}$ | 13.6 ± 1.1 | -7.5 ± 1.1 | -24.0 ± 3.6 | 21.0 ± 1.5 |
| apo-Mb | 7.3 ± 2.1 | -14.0 ± 2.2 | -45.0 ± 7.2 | 21.4 ± 3.1 |

**SI Table 2**. Enthalpy and entropy of activation values derived from Eyring equation. Note that fitting strategy differs from that reported in Sjodt et al. with constrained $k_{fast}$ to 50% of reaction amplitude. Recalculated values are provided for Sjodt et al. Uncertainties are propagated from standard error of the slope and Y-intercept of the Eyring plot (1/T vs. $\ln(k_{fast}/T)$) for each mutant tested.

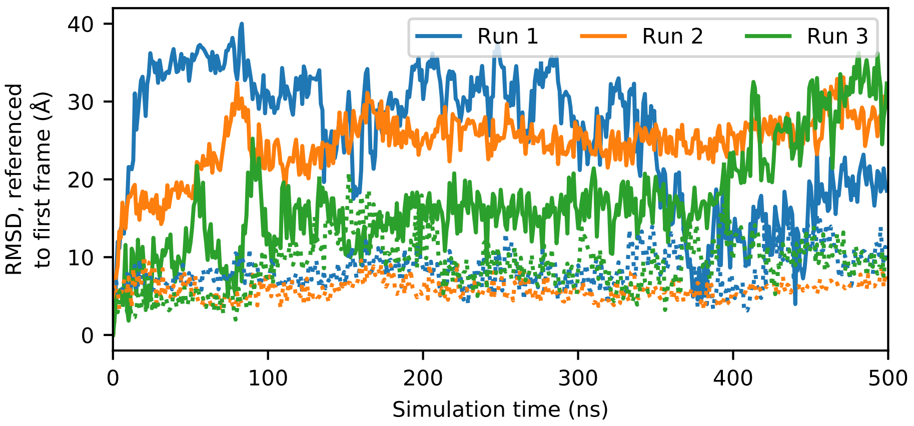


**Figure S1**. RMSD of the N2 domain (solid) and linker-N3 domains (dotted) for the three WT 500ns trajectories after aligning the linking and N3 domains.

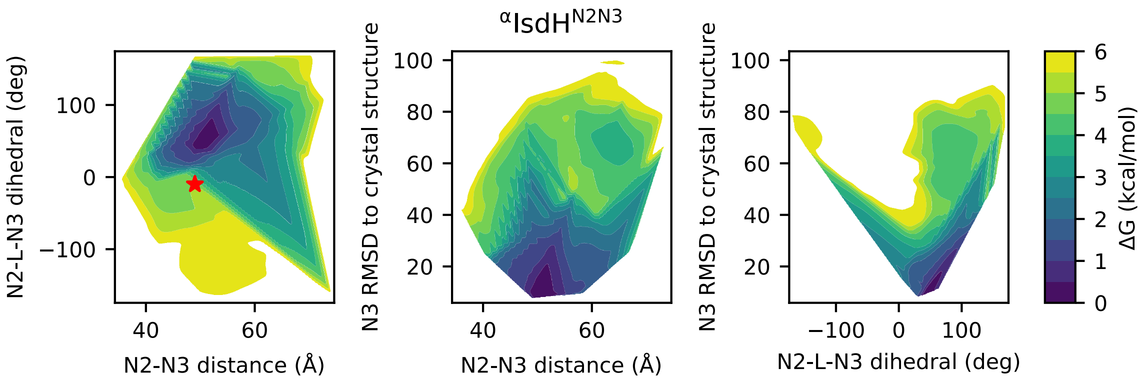


**Figure S2**. Free energy landscapes along the three quantities used to construct the $^{\alpha}$IsdH$^{N2N3}$:Hb Markov model, with the red star representing the configuration seen in the crystal structure.

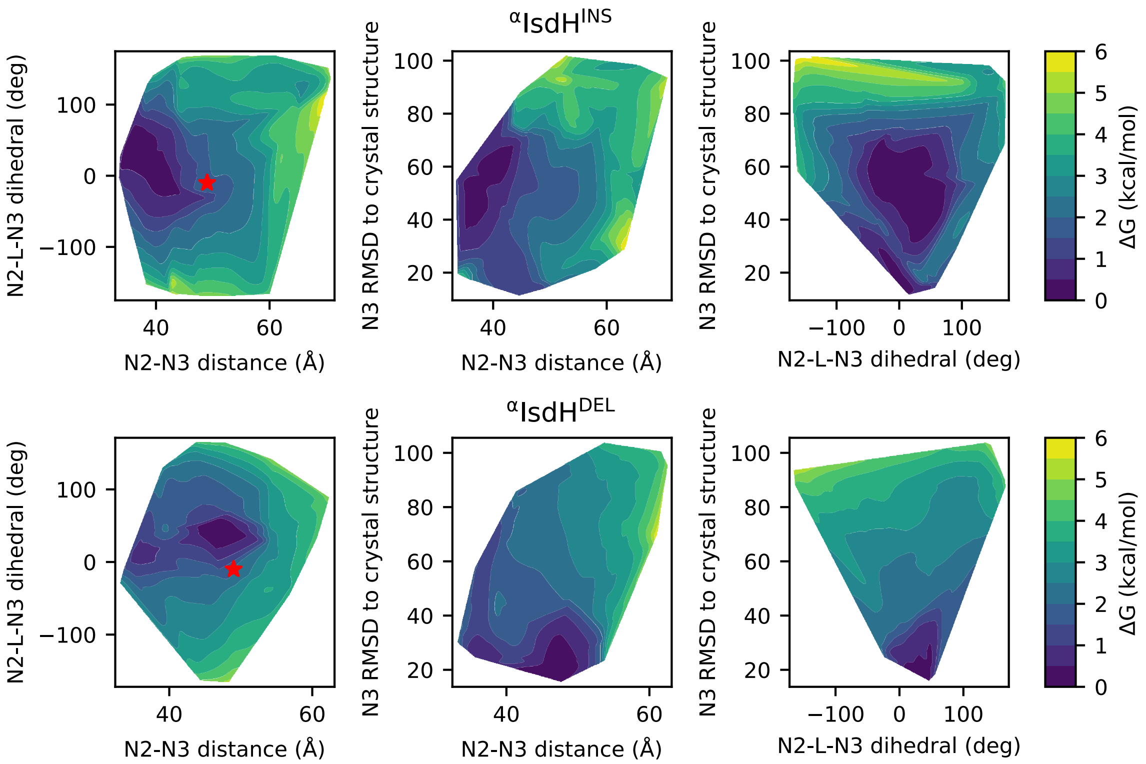


**Figure S3**. Free energy landscapes along the three quantities used to construct the $^{\alpha}$IsdH$^{INS}$:Hb (top) and $^{\alpha}$IsdH$^{DEL}$:Hb (bottom) Markov models, with the red star representing the configuration seen in the crystal structure.
